# Interfering with nucleotide excision by the coronavirus 3’-to-5’ exoribonuclease

**DOI:** 10.1101/2022.08.11.503614

**Authors:** Rukesh Chinthapatla, Mohamad Sotoudegan, Thomas Anderson, Ibrahim M. Moustafa, Kellan T. Passow, Samantha A. Kennelly, Ramkumar Moorthy, David Dulin, Joy Y. Feng, Daniel A. Harki, Robert Kirchdoerfer, Craig E. Cameron, Jamie J. Arnold

## Abstract

Some of the most efficacious antiviral therapeutics are ribonucleos(t)ide analogs. The presence of a 3’-to-5’ proofreading exoribonuclease (ExoN) in coronaviruses diminishes the potency of many ribonucleotide analogs. The ability to interfere with ExoN activity will create new possibilities for control of SARS-CoV-2 infection. ExoN is formed by a 1:1 complex of nsp14 and nsp10 proteins. We have purified and characterized ExoN using a robust, quantitative system that reveals determinants of specificity and efficiency of hydrolysis. Double-stranded RNA is preferred over single-stranded RNA. Nucleotide excision is distributive, with only one or two nucleotides hydrolyzed in a single binding event. The composition of the terminal basepair modulates excision. A stalled SARS-CoV-2 replicase in complex with either correctly or incorrectly terminated products prevents excision, suggesting that a mispaired end is insufficient to displace the replicase. Finally, we have discovered several modifications to the 3’-RNA terminus that interfere with or block ExoN-catalyzed excision. While a 3’-OH facilitates hydrolysis of a nucleotide with a normal ribose configuration, this substituent is not required for a nucleotide with a planar ribose configuration such as that present in the antiviral nucleotide produced by viperin. Design of ExoN-resistant, antiviral ribonucleotides should be feasible.

## Introduction

The emergence of severe acute respiratory syndrome coronavirus 2 (SARS-CoV-2), the causative agent of COVID-19, has led to a global pandemic and caused immeasurable consequences to humankind even more substantial than the incidence of disease and death. While the development of safe and effective vaccines has diminished overall morbidity and mortality, transmission of SARS-CoV-2 continues. Current therapeutics against SARS-CoV-2 infection include polymerase inhibitors: remdesivir and molnupiravir (1,2); and a protease inhibitor: nirmatrelvir, which is boosted with ritonavir (3). All of these therapeutic agents have complications that are tolerable in the midst of the pandemic. However, safer and more effective antiviral agents are needed against multiple targets to support the use of drug cocktails to maximize therapeutic efficacy and minimize the possibility of resistance.

Ribonucleos(t)ide analogs are among the most effective antiviral therapeutics for treatment of RNA virus infections (4-7). This class of compounds is generally administered as the ribonucleoside or ribonucleoside monophosphate prodrug. Cellular kinases then produce the active metabolite, a triphosphorylated ribonucleotide (rNTP) (7). Utilization of the rNTP analog by viral polymerases leads to one or more consequences: chain termination, lethal mutagenesis, backtracking, pausing, and/or recombination, all of which exhibit an antiviral effect (5,6,8-10).

The coronavirus genome is on the order of 30,000 nt, among the largest RNA genome known (11). Because the fidelity of viral RNA polymerases is usually lower than 1 error per 10,000 nt, how coronaviruses resist lethal mutagenesis has been a longstanding question for those studying these viruses (12-14). Efforts initiated in response to emergence of the first SARS-CoV led to the discovery of a 3’-to-5’ proofreading exoribonuclease, termed ExoN (15). ExoN is a member of DEDDh/DEEDh subfamily in the DEED family of exonucleases (13,14,16). These enzymes use a two-metal-ion mechanism for catalysis (13,14,17,18). ExoN is composed of a 1:1 complex of non-structural proteins 14 (60 kDa) and 10 (15 kDa). Nsp14 harbors two catalytic domains: an N-terminal exoribonuclease and a C-terminal methyltransferase (13,14). Nsp10 serves as an accessory factor, stabilizing the active conformation of the exoribonuclease active site and thereby stimulating its activity (19-22).

Consistent with a role of ExoN in proofreading and therefore genome stability, previous studies have shown that genetic inactivation is most often lethal (23). For those coronaviruses, which are viable in the absence of active ExoN, the genomes contain a higher mutational load. Also, these viruses exhibit enhanced sensitivity to some antiviral ribonucleotides (24). Collectively, these observations suggest that inhibitors of ExoN will exhibit antiviral activity and may synergize with ribonucleotide-based inhibitors of the viral polymerase (25). Moreover, an understanding of the determinants of the substrate nucleotide that promote or interfere with excision may guide the design of ExoN-resistant ribonucleotide analogs.

Proofreading DNA exonucleases are generally a subunit of the DNA polymerase holoenzyme and exhibit a preference for mispaired ends (26). This circumstance appears to reflect the preferential partitioning of the mispaired end to the active site of the exonuclease instead of the active site of the polymerase (26,27). How proofreading by ExoN is initiated is not known. Given the dimensions of ExoN and the location of the polymerase active site within the SARS-CoV-2 replication-transcription complex (RTC), an active-site-switching mechanism is unlikely (28,29). It is easy to imagine how an end that cannot be extended by the RTC could become a substrate for repair by ExoN after the RTC dissociates. The ability for nucleotide analogs like remdesivir and molnupiravir to display efficacy in the presence of active ExoN may reflect the inability of these analogs to perturb elongation upon incorporation, with the antiviral activity manifested at the level of the analog-substituted template (30-32).

We have expressed and purified ExoN, and established a robust, quantitative system to study the determinants of the scissile phosphodiester bond and terminal ribonucleotide driving efficient excision by ExoN. ExoN prefers a primed-template-like dsRNA substrate over a ssRNA substrate and cannot access the 3’-terminus at a nick. ExoN only cleaves one or two nucleotides in a single binding event, as expected for a proofreading enzyme (26,27). Cleavage can be completely inhibited by the presence of the phosphorothioate Rp isomer at the scissile phosphodiester bond. A mispaired end does not appear to be highly favored by ExoN relative to a paired end in the absence or presence of a replicating RTC. However, stalling the RTC at the 3’-end of nascent RNA blocks excision regardless of the nature of the basepair. The inability of a mispaired end to promote dissociation of the RTC suggests a role for additional factors in this process. Finally, we identify modifications to the 3’-terminal nucleotide that interfere with excision by ExoN. The most unexpected finding was that the ribose conformation determines whether the 3’-OH is required for efficient turnover. The antiviral nucleotide, ddhCTP, produced by the antiviral protein, viperin, is readily excised by ExoN. This molecule lacks a 3’-OH but also lacks sugar pucker because of the presence of a double bond forcing the ribose into a planar conformation. This observation is consistent with repair of ddhC-terminated RNA as a major driver for acquisition ExoN and evolution of its substrate specificity.

## Materials and Methods

### Materials

DNA oligonucleotides and dsDNA fragments, GBlocks, were from Integrated DNA Technologies. RNA oligonucleotides were either from Horizon Discovery Ltd. (Dharmacon) or Integrated DNA Technologies. Restriction enzymes and T4 PNK (3’-phosphatase minus) were from New England Biolabs. IN-FUSION HD enzyme was from TakaraBio. Phusion DNA polymerase and T4 polynucleotide kinase were from ThermoFisher. pBirACm plasmid DNA was from Avidity. Streptactin XT 4F High-Capacity resin was from IBA Life Sciences. [γ-^32^P]ATP (6,000 Ci/mmol) and [α-^32^P]ATP (3,000 Ci/mmol) were from Perkin Elmer. Nucleoside 5’-triphosphates (ultrapure solutions) were from Cytiva. pET16b-RtcA-NTerm-His expression plasmid was provided by Stewart Shuman (Sloan Kettering) (33). Cytidine 5’-O-(1-thiotriphosphate) and 3’-deoxycytidine 5’-triphosphate were from TriLink. Adenosine 5’-O-(1-thiotriphosphate) (Sp isomer) was from BioLog. Remdesivir-terminated RNA was synthesized by both Dharmacon and the Harki lab as previously described (34). Remdesivir was provided to Dharmacon by Gilead Sciences, and this was chemically converted to the phosphoramidite to be synthetically incorporated into RNA. ddhCTP was synthesized by the Harki lab as previously described (35); 2’-C-methylcytidine 5’-triphosphate was provided by Gilead Sciences. All other reagents were of the highest grade available from MilliporeSigma, VWR, or Fisher Scientific.

### Construction of modified pSUMO vectors containing AviTag

The pSUMO system allows for production of SUMO fusion proteins containing an amino-terminal affinity tag fused to SUMO that can be purified by affinity chromatography and subsequently processed by the SUMO protease, Ulp1 (36). After cleavage this will produce an authentic untagged protein target of interest. The pSUMO vector (LifeSensors) (36) was modified such that the coding sequence for the six-histidine tag was replaced with a DNA sequence coding for an AviTag codon optimized for bacterial expression (37,38). The AviTag is a short 15 amino acid sequence (GLNDIFEAQKIEWHE) that can be specifically biotinylated on the lysine residue by the biotin ligase, BirA (37,38). The construct contains two tags separated by a short linker to increase the affinity to the chromatography resin during purification. The biotinylated AviTag allows affinity purification of the fusion protein to be isolated using streptavidin or Strep-Tactin resins. The DNA sequences (GBlocks encoding the tags) were cloned into pSUMO using XbaI and SalI by IN-FUSION. The final constructs (pAviTag_SUMO) were confirmed by sanger sequencing performed by Genewiz.

### Codon-optimized sequence for AviTag

5’ATGGGACTAAATGATATATTTGAAGCTCAAAAGATCGAGTGGCACGAGGGTGGTGGCAGCGGTGGCGGCTCCGGCGGTAGCGGCCTGAACGACATCTTCGAGGCGCAGAAAATTGAATGGCATGAAGGTGGCTCTAGCGGTGGT3’ (amino acid sequence: MGLNDIFEAQKIEWHEGGGSGGGSGGSGLNDIFEAQKIEWHEGGSSGG)

### Construction of SARS-CoV-2 nsp10 and nsp14 bacterial expression plasmids

The SARS-CoV-2 nsp10 and nsp14 genes were codon optimized for expression in *E. coli* and obtained from Genescript. The amino acid sequences for nsp10 and nsp14 were derived from SARS-CoV-2 isolate 2019-nCoV/USA-WA1/2020 (GenBank MN985325.1). The genes were amplified by PCR using Phusion DNA polymerase. The nsp10 gene was amplified using the synthetic nsp10 gene as template and DNA oligonucleotides (5’-GAACAGATTGGAGGTGCCGGGAATGCTACGGAA-3’ and 5’-CCGCAAGCTTGTCGACTTATCATTGAAGCATAGGTTCACGCAA-3’). The nsp14 gene was amplified using the synthetic nsp14 gene as template and DNA oligonucleotides (5’-GAACAGATTGGAGGTGCGGAGAACGTTACAGGT and 5’-CCGCAAGCTTGTCGACTTATCATTGAAGGCCAGTAAACGTATTCCA-3’). The PCR products were gel purified and cloned into either the pAviTag-pSUMO bacterial expression plasmids using BsaI and SalI. The final constructs were confirmed by sanger sequencing performed by Genewiz. To construct a catalytically inactive nsp14, the WT nsp14 expression plasmid was modified such that D90 and E92 were both changed to alanine. This was performed using Quickchange mutagenesis using the WT nsp14 expression plasmid as template and DNA oligonucleotides (5’-GCGTGGATTGGTTTTGCTGTTGCGGGTTGCCACGCGACCCGT-3’ and 5’-ACGGGTCGCGTGGCAACCCGCAACAGCAAAACCAATCCACGC-3’). The final construct was confirmed by sanger sequencing performed by Genewiz.

### Codon optimized sequence for SARS-CoV-2 nsp10

5’GCCGGGAATGCTACGGAAGTTCCAGCTAACTCGACCGTTCTTAGCTTTTGTGCTTTTGCAGTCGATGCAGCGAAAGCGTATAAGGACTATCTGGCGTCAGGGGGACAACCCATTACTAACTGTGTCAAGATGCTGTGTACCCATACCGGCACGGGTCAAGCGATTACTGTTACACCAGAAGCTAACATGGACCAGGAATCTTTTGGTGGTGCCAGTTGCTGCTTGTACTGCCGCTGTCATATCGATCACCCCAATCCAAAAGGTTTCTGCGATCTGAAGGGAAAATACGTGCAAATCCCCACCACTTGTGCTAATGACCCGGTCGGATTTACGCTGAAGAACACCGTTTGTACTGTTTGCGGGATGTGGAAAGGGTATGGGTGTTCTTGCGACCAGTTGCGTGAACCTATGCTTC AA3’

### Codon optimized sequence for SARS-CoV-2 nsp14

5’GCGGAGAACGTTACAGGTTTATTTAAGGATTGCTCTAAAGTAATTACCGGCCTGCACCCAACGCAGGCACCAACTCATCTTAGCGTGGATACAAAATTTAAGACAGAAGGACTGTGTGTGGACATTCCTGGCATCCCAAAGGACATGACATACCGCCGTTTGATCTCCATGATGGGGTTCAAAATGAACTACCAGGTAAACGGATACCCTAATATGTTCATTACACGTGAGGAGGCGATTCGTCATGTCCGCGCCTGGATCGGATTCGACGTAGAAGGTTGCCACGCCACCCGTGAGGCTGTGGGGACGAACTTACCCCTTCAGCTTGGCTTCTCAACTGGGGTAAACTTGGTGGCCGTCCCGACAGGGTATGTTGACACTCCTAATAACACTGATTTCTCGCGTGTATCTGCAAAGCCACCACCAGGGGACCAGTTCAAACACCTGATCCCCCTGATGTATAAGGGTCTTCCTTGGAATGTGGTCCGTATTAAAATCGTCCAGATGCTGTCAGACACCCTTAAGAATCTGTCAGATCGTGTGGTATTTGTATTGTGGGCGCACGGATTCGAGTTAACAAGCATGAAATATTTTGTGAAAATTGGCCCCGAACGCACATGCTGCTTATGCGATCGTCGCGCTACTTGCTTTAGTACTGCTTCAGACACTTATGCCTGCTGGCACCACTCTATTGGATTTGACTACGTGTATAACCCATTCATGATTGATGTCCAGCAGTGGGGCTTCACCGGGAACTTGCAGTCCAACCATGACCTTTATTGTCAGGTTCACGGAAATGCCCACGTGGCAAGCTGCGACGCGATTATGACACGCTGTCTGGCGGTACATGAGTGCTTTGTAAAGCGTGTCGATTGGACCATCGAGTATCCAATCATTGGAGACGAACTTAAGATCAATGCCGCATGCCGTAAAGTTCAACACATGGTAGTAAAGGCCGCCCTTCTTGCGGATAAGTTTCCGGTTCTGCATGACATTGGCAACCCTAAGGCGATTAAGTGTGTCCCGCAGGCGGATGTCGAATGGAAATTCTATGACGCGCAACCCTGCTCGGATAAAGCATATAAAATCGAAGAGCTGTTTTATTCATACGCTACGCATTCCGACAAGTTTACAGATGGCGTTTGTCTTTTTTGGAATTGTAACGTTGATCGCTACCCGGCGAACTCAATCGTTTGCCGCTTTGACACACGTGTGCTGTCTAACTTGAACTTGCCTGGTTGCGATGGAGGCTCGTTGTATGTTAATAAACATGCGTTTCATACCCCCGCCTTCGACAAGTCCGCTTTCGTAAACCTGAAGCAGTTGCCATTTTTCTACTATAGCGACTCACCGTGCGAGTCCCACGGTAAGCAAGTAGTGTCTGACATTGATTATGTACCTTTAAAAAGTGCTACCTGCATCACCCGTTGCAACTTGGGCGGAGCGGTTTGCCGCCACCA TGCGAACGAATATCGCTTATACCTTGATGCCTATAATATGATGATTAGCGCGGGATTTAGCCTTTGGGTTTATAAACAGTTCGATACTTATAACCTGTGGAATACGTTTACTCGCCTTCAA3’

### Expression and purification of SARS-CoV-2 nsp10

*E. coli* BL21(DE3)pBirACm competent cells were transformed with the pAviTag_SUMO-SARS-CoV-2-nsp10 plasmid for protein expression. BL21(DE3)pBirACm cells containing the pAviTag_SUMO-SARS-CoV-2-nsp10 plasmid were grown in 100 mL of media (NZCYM) supplemented with kanamycin (K25, 25 µg/mL) and chloramphenicol (C20, 20 µg/mL) at 37 °C until an OD_600_ of 1.0 was reached. This culture was then used to inoculate 4L of K25,C20 media to an OD_600_ = 0.1. Biotin (25 mM in 500 mM Bicine pH 8.0) was added to the media to a final concentration of 50 µM. The cells were grown at 37 °C to an OD_600_ of 0.8 to 1.0, cooled to 25 °C and then IPTG (500 µM) was added to induce protein expression. Cultures were then grown for an additional 4 h at 25 °C. Cells were harvested by centrifugation (6000 x *g*, 10 min) and the cell pellet was washed once in 200 mL of TE buffer (10 mM Tris, 1 mM EDTA), centrifuged again, and the cell paste weighed. The cells were then frozen and stored at -80 °C until used. Frozen cell pellets were thawed on ice and suspended in lysis buffer (25 mM HEPES pH 7.5, 500 mM NaCl, 2 mM TCEP, 20% glycerol, 1.4 μg/mL leupeptin, 1.0 μg/mL pepstatin A and two Roche EDTA-free protease tablet per 5 g cell pellet), with 5 mL of lysis buffer per 1 gram of cells. The cell suspension was lysed by passing through a French press (SLM-AMINCO) at 15,000 psi. After lysis, phenylmethylsulfonylfluoride (PMSF) and NP-40 were added to a final concentration of 1 mM and 0.1% (v/v), respectively. While stirring the lysate, polyethylenimine (PEI) was slowly added to a final concentration of 0.25% (v/v) to precipitate nucleic acids from cell extracts. The lysate was stirred for an additional 30 min at 4 °C after the last addition of PEI, and then centrifuged at 75,000 x g for 30 min at 4 °C. The PEI supernatant was then loaded onto a Strep-Tactin XT 4F HC resin (IBA Life Sciences) at a flow rate of 1 mL/min (approximately 1 mL bed volume per 100 mg total protein) equilibrated with buffer A (25 mM HEPES, pH 7.5, 500 mM NaCl, 2 mM TCEP, 20% glycerol). After loading, the column was washed with twenty column volumes of buffer A. The resin was then suspended in two column volumes of buffer A and Ulp1 (5 µg per 1 mL bed volume) was added with the resin overnight at 4 °C to cleave the SUMO-nsp10 fusion protein. The column was then washed with 5 column volumes of buffer A and fractions were collected and assayed for purity by SDS-PAGE. The protein concentration was determined by measuring the absorbance at 280 nm by using a Nanodrop spectrophotometer and using a calculated molar extinction coefficient of 13,700 M^-1^ cm^-1^. Purified, concentrated protein was aliquoted and frozen at -80 °C until use. Typical nsp10 yields were 1 mg/1 g of *E. coli* cells.

### Expression and purification of SARS-CoV-2 nsp14

*E. coli* BL21(DE3)pBirACm competent cells were transformed with the pAviTag_SUMO-SARS-CoV-2-nsp14 plasmid for protein expression. BL21(DE3)pBirACm cells containing the pAviTag_SUMO-SARS-CoV-2-nsp14 plasmid were grown in 100 mL of media (NZCYM) supplemented with kanamycin (K25, 25 µg/mL) and chloramphenicol (C20, 20 µg/mL) at 37 °C until an OD600 of 1.0 was reached. This culture was then used to inoculate 4L of K25,C20 media to an OD600 = 0.1. Biotin (25 mM in 500 mM Bicine pH 8.0) was added to the media to a final concentration of 50 µM. The cells were grown at 37 °C to an OD600 of 0.8 to 1.0, cooled to 25 °C and then IPTG (500 µM) was added to induce protein expression. Cultures were then grown for an additional 4 h at 25 °C. Cells were harvested by centrifugation (6000 x *g*, 10 min) and the cell pellet was washed once in 200 mL of TE buffer (10 mM Tris, 1 mM EDTA), centrifuged again, and the cell paste weighed. The cells were then frozen and stored at -80 °C until used. Frozen cell pellets were thawed on ice and suspended in lysis buffer (25 mM HEPES pH 7.5, 500 mM NaCl, 2 mM TCEP, 20% glycerol, 1.4 μg/mL leupeptin, 1.0 μg/mL pepstatin A and two Roche EDTA-free protease tablet per 5 g cell pellet), with 5 mL of lysis buffer per 1 gram of cells. The cell suspension was lysed by passing through a French press (SLM-AMINCO) at 15,000 psi. After lysis, phenylmethylsulfonylfluoride (PMSF) and NP-40 were added to a final concentration of 1 mM and 0.1% (v/v), respectively. While stirring the lysate, polyethylenimine (PEI) was slowly added to a final concentration of 0.25% (v/v) to precipitate nucleic acids from cell extracts. The lysate was stirred for an additional 30 min at 4 °C after the last addition of PEI, and then centrifuged at 75,000 x g for 30 min at 4 °C. The PEI supernatant was then loaded onto a Strep-Tactin XT 4F HC resin (IBA Life Sciences) at a flow rate of 1 mL/min (approximately 1 mL bed volume per 100 mg total protein) equilibrated with buffer A (25 mM HEPES, pH 7.5, 500 mM NaCl, 2 mM TCEP, 20% glycerol). After loading, the column was washed with twenty column volumes of buffer A. The resin was then suspended in two column volumes of buffer A and Ulp1 (5 µg per 1 mL bed volume) was added with the resin overnight at 4 °C to cleave the SUMO-nsp14 fusion protein. The column was then washed with 5 column volumes of buffer A and fractions were collected and assayed for purity by SDS-PAGE. The protein concentration was determined by measuring the absorbance at 280 nm by using a Nanodrop spectrophotometer and using a calculated molar extinction coefficient of 93,625 M^-1^ cm^-1^. Purified, concentrated protein was aliquoted and frozen at -80 °C until use. Typical nsp14 yields were 0.2 mg/1 g of *E. coli* cells. The catalytically inactive nsp14 D90A E92A was expressed and purified using the exact same conditions as WT nsp14.

### SARS CoV-2 nsp7, nsp8, and nsp12 recombinant proteins

Expression and purification of SARS CoV-2 nsp7, nsp8 and nsp12 are described in detail in (8).

### Expression and purification of RtcA

*E. coli* Rosetta(DE3) competent cells were transformed with the pET16b-RtcA-NTerm-His plasmid for protein expression. *E. coli* Rosetta(DE3) cells containing the pET16b-RtcA-NTerm-His plasmid were grown in 100 mL of A300,C60-supplemented ZYP-5052 auto-induction media at 37 °C (39,40). The cells were grown at 37 °C to an OD_600_of 0.8 to 1.0, cooled to 15 °C and then grown for 36-44 h. After ∿40 h at 15 °C the OD_600_reached ∿10– 15. Cells were harvested by centrifugation (6000 x *g*, 10 min) and the cell pellet was washed once in 200 mL of TE buffer (10 mM Tris, 1 mM EDTA), centrifuged again, and the cell paste weighed. The cells were then frozen and stored at -80 °C until used. Frozen cell pellets were thawed on ice and suspended in lysis buffer (25 mM HEPES pH 7.5, 500 mM NaCl, 2 mM TCEP, 20% glycerol, 5 mM imidazole, 1.4 μg/mL leupeptin, 1.0 μg/mL pepstatin A and two Roche EDTA-free protease tablet per 5 g cell pellet), with 5 mL of lysis buffer per 1 gram of cells. The cell suspension was lysed by passing through a French press (SLM-AMINCO) at 15,000 psi. After lysis, phenylmethylsulfonylfluoride (PMSF) and NP-40 were added to a final concentration of 1 mM and 0.1% (v/v), respectively. The lysate was centrifuged at 75,000 x g for 30 min at 4 °C. The lysate was then loaded onto Qiagen Ni-Spin columns (a total of 5 mL per one Ni spin column). After loading, the column was washed with two 0.5 mL volumes of buffer B (25 mM HEPES, pH 7.5, 500 mM NaCl, 2 mM TCEP, 20% glycerol) with 5 mM imidazole, then washed with two 0.5 mL volumes of buffer B with 50 mM imidazole and then eluted in four 0.1 mL volumes of buffer B with 500 mM imidazole. Elution fractions were collected and assayed for purity by SDS-PAGE. The protein concentration was determined by measuring the absorbance at 280 nm by using a Nanodrop spectrophotometer and using a calculated molar extinction coefficient of 11,585 M^-1^ cm^-1^. Purified, concentrated protein was aliquoted and frozen at -80 °C until use. RtcA yields were 0.5 mg/2 g of *E. coli* cells.

### 5’-^32^P-labeling of RNA substrates

RNA oligonucleotides were end-labeled by using [*γ*-^32^P]ATP and T4 polynucleotide kinase. Reaction mixtures, with a typical volume of 50 *μ*L, contained 0.5 *μ*M [*γ*-^32^P]ATP, 10 *μ*M RNA oligonucleotide, 1× kinase buffer, and 0.4 unit/*μ*L T4 polynucleotide kinase. Reaction mixtures were incubated at 37 °C for 60 min and then held at 65 °C for 5 min to heat inactivate T4 PNK. For RNAs containing a 3’phosphate T4 PNK (minus 3’phosphatase) was used using the same reaction conditions.

### Cyclization reactions to produce 2’,3’-cyclic phosphate containing RNAs

Reactions contained 25 mM HEPES pH 7.5, 10 mM MgCl2, 10 mM TCEP, 50 mM NaCl, 100 µM ATP, 1 µM ^32^P-labeled RNA (3’-phosphate termini) and 5 µM RtcA. Reactions were performed at 37 °C for 30 min. Reactions were quenched by addition of EDTA to a final concentration of 10 mM and placed at 65 °C to heat inactivate RtcA enzyme.

### Annealing of dsRNA substrates

dsRNA substrates were produced by annealing 10 μM RNA oligonucleotides in T10E1 [10 mM Tris pH 8.0 and 1 mM EDTA] and 50 mM NaCl in a Progene Thermocycler (Techne). Annealing reaction mixtures were heated to 90 °C for 1 min and slowly cooled (5 °C/min) to 10 °C. Specific scaffolds are described in the figure legends.

### SARS-CoV-2 nsp10/nsp14-catalyzed exoribonuclease assays

Reactions contained 25 mM HEPES pH 7.5, 2 mM MgCl2, 1 mM TCEP, 10 mM KCl, and 50 mM NaCl and were performed at 30 °C. Reactions were quenched by addition of EDTA to a final concentration of 25 mM. Specific concentrations of RNA substrate, ExoN, along with any deviations from the above, are indicated in the appropriate figure legend. Typical concentrations for RNA substrate and enzyme were between 0.01 to 2 µM. Enzymes were diluted immediately prior to use in 25 mM HEPES, pH 7.5, 500 mM NaCl, 2 mM TCEP, and 20% glycerol. SARS-CoV-2 nsp10 was pre-mixed with SARS-CoV-2 nsp14 on ice in 25 mM HEPES, pH 7.5, 500 mM NaCl, 2 mM TCEP, and 20% glycerol for 5 min prior to initiating the reaction with ExoN. The volume of enzyme added to any reaction was always less than or equal to one-tenth the total volume. The ExoN concentration refers to the nsp14 concentration and the ratio of nsp14 to nsp10 is also indicated in cases where nsp10 was in excess of nsp14. For example, 0.1 µM ExoN (1:5) refers to final concentrations of nsp14 of 0.1 µM and nsp10 at 0.5 µM. Products were resolved from substrates by denaturing PAGE.

### Incorporation of modified nucleoside triphosphates into dsRNA substrates prior to challenge for removal by ExoN

Reactions contained 25 mM HEPES pH 7.5, 5 mM MgCl2, 1 mM TCEP and 50 mM NaCl and were performed at 30 °C. Human mitochondrial RNA polymerase, POLRMT, was used to incorporate modified nucleoside triphosphates. POLRMT was expressed and purified as described previously (41). 1 µM POLRMT was mixed with 1 µM ^32^P-labeled dsRNA nucleic acid scaffold (primed-template) in the presence of 10 µM ATP, 10 µM UTP, and either 10 µM CTP or 100 µM of the modified nucleoside 5’-triphosphate. The RNA primer was 9P (5’-CCGGGCGGC-3’) and RNA template was 21T (**Table 1**). This primer-template pair allows ATP to be incorporated at the n+1 position, UTP at n+2 and the modified nucleoside 5’-triphosphate to be incorporated at the n+3 position. Reactions were allowed to proceed for 30 min to allow complete extension to n+3 at which point 0.1 µM ExoN (1:5) was added to the reaction. Reactions were quenched at various times by addition of 25 mM EDTA. Products were resolved from substrates by denaturing PAGE.

**Table 1.**
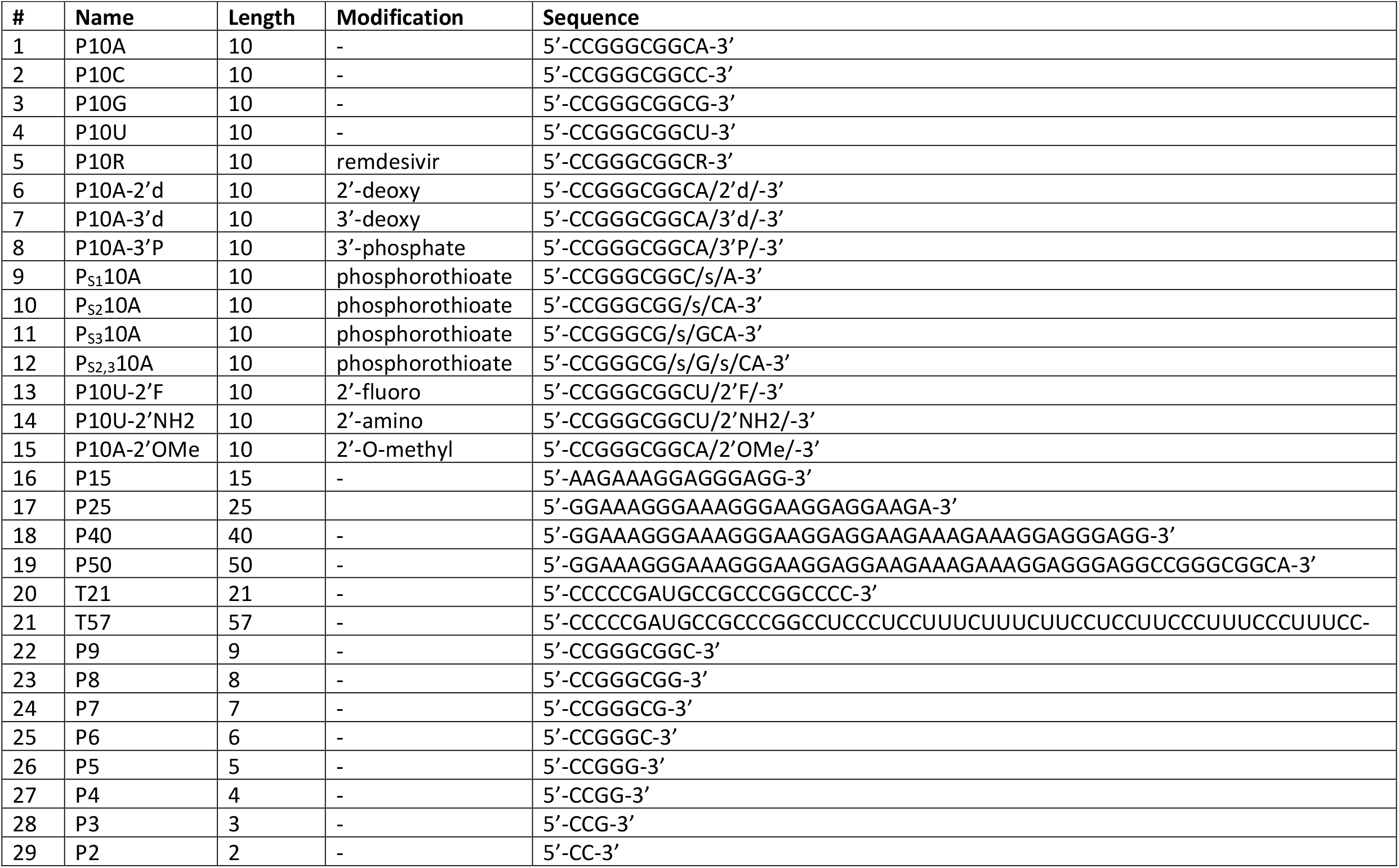
RNA Oligonucleotides. RNA substrates used in this study. The name, length, type of modification (if any), and sequence are indicated.

### Incorporation of correct and incorrect nucleotides by SARS-CoV-2 replicase into dsRNA substrates prior to challenge for removal by ExoN

Reactions contained 25 mM HEPES pH 7.5, 2 mM MgCl2, 1 mM TCEP and 50 mM NaCl and were performed at 30 °C. Reactions contained 0.1 µM nsp12, 0.3 µM nsp7, 0.3 µM nsp8, 0.1 µM ExoN (1:5), 0.1 µM dsRNA primed/template (9P/40AG/57T), 1 µM ATP, 0.1 µCi/µL [α-^32^P]ATP, 1 µM UTP, and either 10 µM CTP or 100 µM of the modified nucleoside 5’-triphosphate. Initially, nsp12/7/8 was incubated with dsRNA substrate and ATP in the absence or presence of UTP for 60 min to form n+1 or n+2 (with UTP) product, at which point either CTP, 3’-dCTP, ddhCTP, 2’-C-Me-CTP was added to promote further extension. After 2 min, ExoN was added, and the reaction was quenched at various times by the addition of EDTA to 25 mM.

### Denaturing PAGE analysis of exonuclease-catalyzed reaction products

An equal volume of loading buffer (85% formamide, 0.025% bromophenol blue and 0.025% xylene cyanol) was added to quenched reaction mixtures and heated to 90 °C for 5 min prior to loading 5 µL on a denaturing either 15% or 23% polyacrylamide gel containing 1X TBE (89 mM Tris base, 89 mM boric acid, and 2 mM EDTA) and 7 M urea. For reactions that contained dsRNA substrates an excess (50-fold) of unlabeled RNA oligonucleotide (trap strand) that is the exact same sequence to the ^32^P-labeled RNA oligonucleotide in the reaction was present in the loading buffer to ensure complete separation and release of the ^32^P-lableled RNA oligonucleotide prior to gel electrophoresis (42). This procedure allows efficient strand separation of ^32^P-labeled RNA oligos that are in the presence of their RNA complements (42). Electrophoresis was performed in 1x TBE at 90 W. Gels were visualized by using a PhosphorImager (GE) and quantified by using ImageQuant TL software (GE).

### Data analysis

All gels shown are representative, single experiments that have been performed at least three to four individual times to define the concentration or time range shown with similar results. In all cases, values for parameters measured during individual trials were within the limits of the error reported for the final experiments. Data were fit by either linear or nonlinear regression using the program GraphPad Prism v7.03 (GraphPad Software Inc.).

## Results

### Expression and purification of SARS-CoV-2 ExoN: nsp10 and nsp14

Many proteins and enzymes encoded by SARS-CoV-2 contain zinc ions coordinated by side chains of histidine and/or cysteine residues, including the nsp10 and nsp14 subunits of the 3’-to-5’ exoribonuclease complex (ExoN) (11,13,14,19). For the nsp12 gene-encoded RdRp, the natural ligand has been proposed to be a four-iron, four-sulfur cluster (43). To avoid the potential for displacement of metal ions during protein purification that were incorporated during protein expression, we used the AviTag-SUMO system, which avoids metal-affinity chromatography (**Fig. S1A**)(37,38). The AviTag is a 15 amino acid sequence (GLNDIFEAQKIEWHE) that is biotinylated on the lysine residue by biotin ligase, BirA, co-expressed with the fusion protein (37,38). The biotinylated AviTag-SUMO fusion protein binds to streptavidin or streptactin resins (**Fig. S1B**) (37,38). The protein of interest is released from the resin by cleavage at the carboxyl terminus of SUMO by the Ulp1 protease (**Figs. S1C-F**) (38). Purified proteins used in this study are shown in **Fig. 1A**. We also purified a catalytically inactive derivative of nsp14, referred to as MUT (**Fig. S1F**).

**Figure 1.**
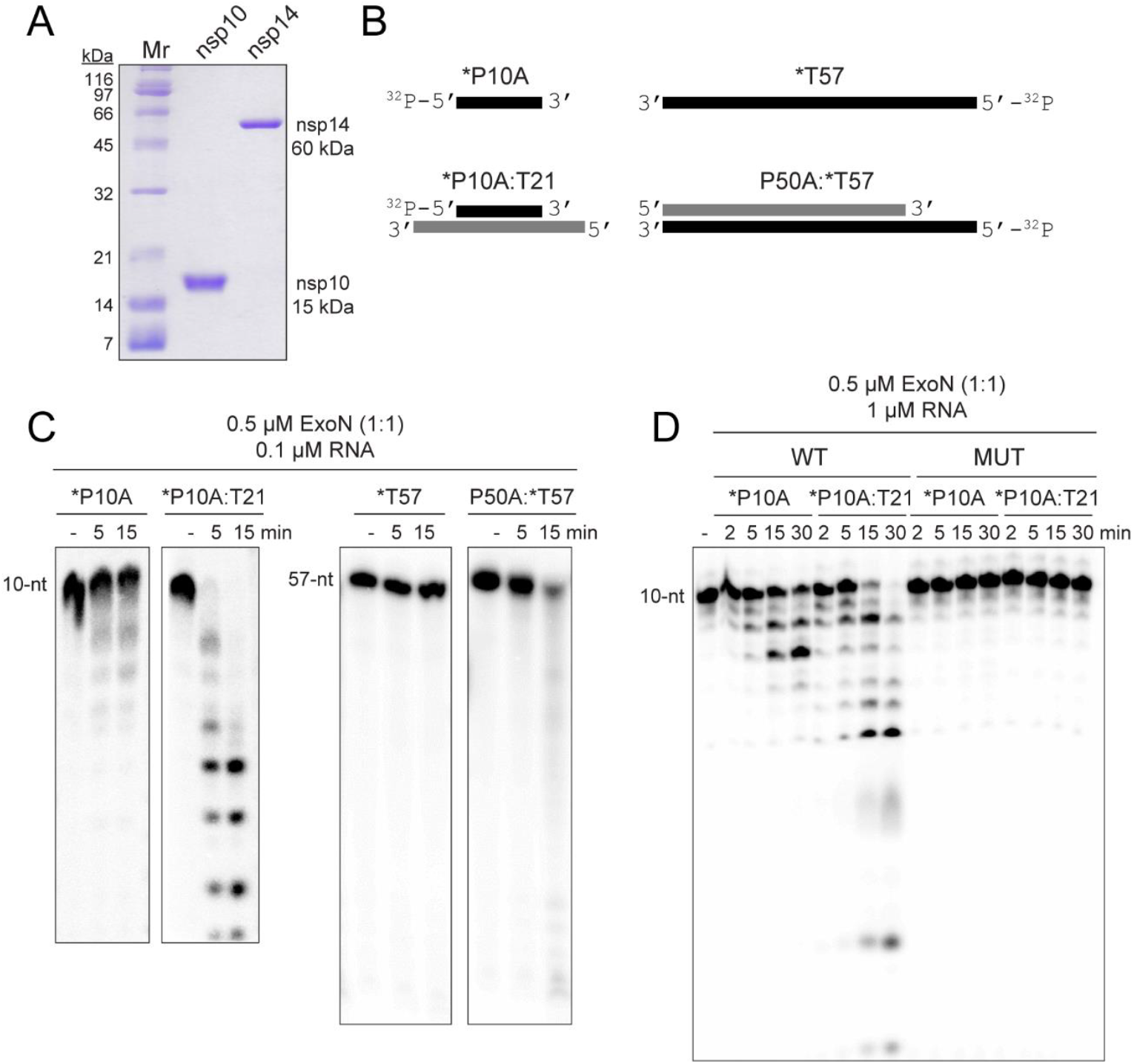
ExoN prefers primed-template-like dsRNA substrates. (**A**) Purified SARS-CoV-2 nsp14 (60 kDa) and nsp10 (15 kDa) proteins used in this study. Proteins (1 µg) were resolved on a 15% polyacrylamide gel containing SDS and stained with Coomassie. Broad-range molecular weight markers (Mr) and corresponding molecular weights are indicated. (**B**) Schematic of ssRNA and dsRNA substrates used to measure exoribonuclease activity. RNA sequences are provided in **Table 1** and/or **Fig. S2**. RNAs were labeled on the 5’-end with ^32^P, also indicated by an asterisk (*). RNAs are designated as primers (P) or templates (T). The numbers indicate the length of the RNA. When a base is designated, this reflects the 3’-terminal nucleotide. (**C**) Evaluation of ExoN-catalyzed hydrolysis of RNA. Reaction products were resolved by denaturing PAGE and visualized by phosphorimaging. Reactions contained 0.1 µM ExoN (1:1 nsp14:nsp10) and 0.1 µM of the indicated RNA, were incubated at 30 °C for the indicated time, then quenched by addition of EDTA. In both cases, primed-template-like dsRNA substrates were cleaved more efficiently than ssRNA substrates. (**D**) Exoribonuclease activity is dependent on the nsp14 active site. Residues of nsp14 required for catalysis were changed as follows: D90A, E92A; this derivative is referred to as MUT. Reactions contained 0.1 µM ExoN (1:1 nsp14(WT or MUT):nsp10) and 1 µM RNA and were run as described above. MUT did not exhibit any detectable exoribonuclease activity.

### ExoN prefers a dsRNA substrate containing a recessed 3’-end, a “primed-template”

We used ssRNA and dsRNA substrates of various lengths in the assay (**Fig. 1B, Table 1, Fig. S2**). We chose these substrates because they can also be used as primers (P) and templates (T) for the replication-transcription complex (RTC). We use denaturing polyacrylamide gel electrophoresis followed by phosphorimaging to monitor the hydrolysis of the ^32^P-labeled RNA strand (indicated by an asterisk) in the absence (ssRNA) or presence of an annealed RNA strand (dsRNA). The concentration of ExoN and nsp14:nsp10 stoichiometry used was selected to reflect conditions used by others (19,20,23,28,44-51), to facilitate a comparison of the results obtained here to those. Our studies demonstrated that ssRNA was not a good substrate for ExoN (*P10A and *T57 in **Fig. 1C**). The 3’-end of a blunt-ended duplex was also a poor substrate for ExoN (P50A:*T57 in **Fig. 1C**). dsRNA with a recessed 3’-end was the most active substrate (*P10A:T21 in **Fig. 1C**). Comparable results were obtained with P50A and T57 RNAs (**Fig. S3**). Mutagenesis of the nsp14 ligands required for Mg^2+^-dependent catalysis (D90A, E92A) inactivated ExoN (compare MUT to WT in **Fig. 1D**) (13-16). Therefore, none of the activity measured here derived from a contaminating ribonuclease.

### Products of the nsp15 endonuclease are not substrates for ExoN

Coronaviruses also encode an uridylate-specific, ssRNA or dsRNA endonuclease, referred to as NendoU and is encoded by the nsp15-coding sequence of the SARS-CoV-2 genome (52,53). While the activity on dsRNA has been suggested to clear dsRNA that would otherwise activate intrinsic antiviral defenses (52,54,55), the possibility existed that this enzyme could contribute to a post-transcriptional, mismatch-repair mechanism. For example, nsp15 might also cleave the phosphodiester backbone at or near a mismatch. If ExoN could cleave at a nick or at a 2’-3’ cyclic-phosphate or 3’-phosphate terminus produced by nsp15, then such a repair mechanism might exist.

To test this possibility, we assembled substrates in which the P50A RNA was fragmented into three segments: P-1, P-2, and P-3 in the 5’-to-3’ direction, respectively (**Fig. 2A** and **Fig. S2A**). We evaluated three conditions: I, where P-3 was labeled; II, where P-3 was omitted and P-2 was labeled; and III, where P-2 was labeled (**Fig. 2A**). The labeled, 3’-terminal P-3 primer was cleaved efficiently (I in **Fig. 2B**). Similarly, P-2 was cleaved efficiently when present as the 3’-terminal RNA (II in **Fig. 2B**). However, P-2 was not cleaved when embedded between P-1 and P-3 (III in **Fig. 2B**), suggesting that ExoN is incapable of initiating hydrolysis from the 3’-OH at a nick.

**Figure 2.**
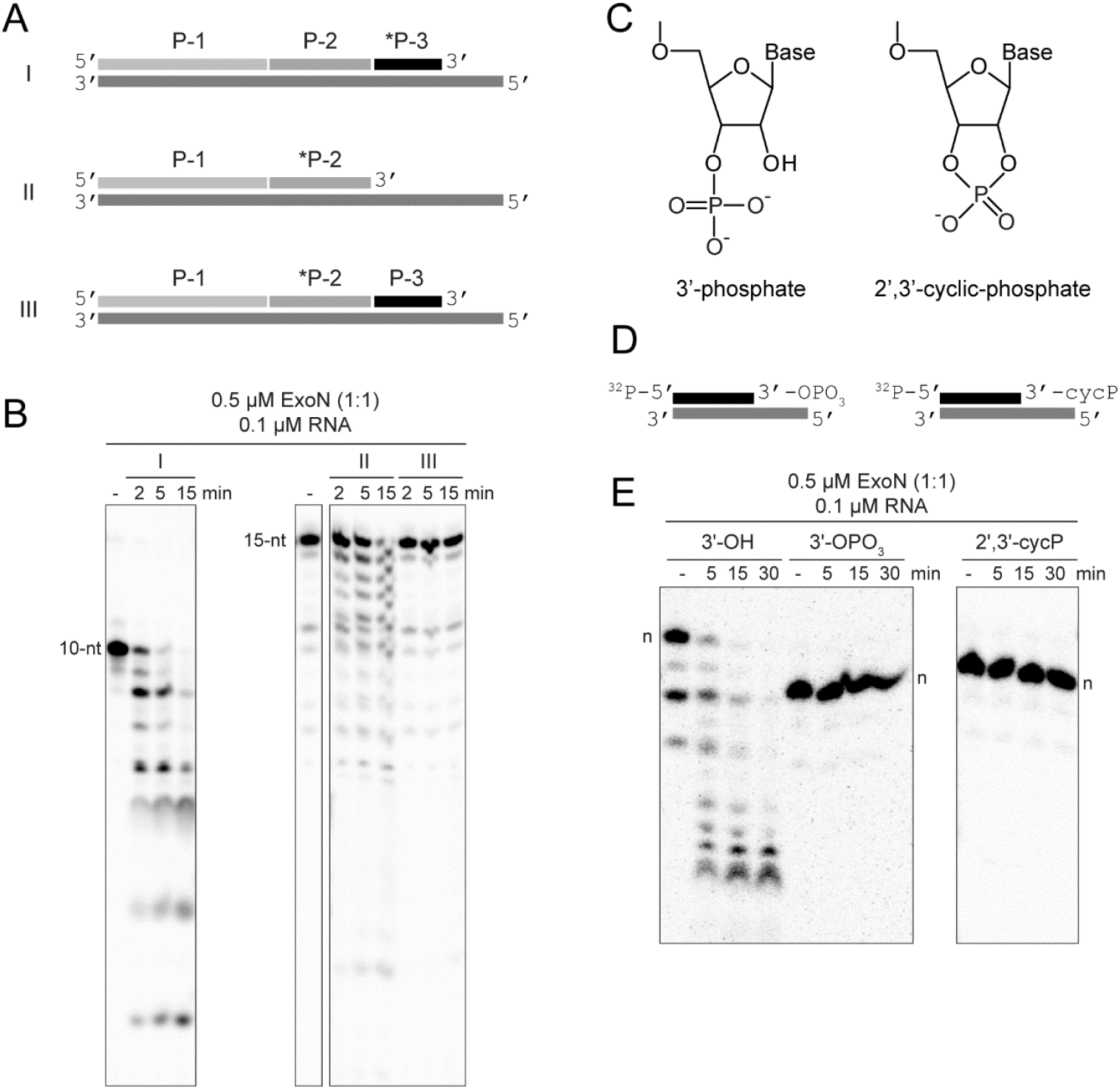
Products of nsp15-catalyzed endonucleolytic cleavage are not substrates for ExoN. (**A**) Schematic of dsRNA substrates used. Both substrates I and II contain ^32^P-labeled RNAs (*P-3 and *P-2) annealed to the template such that both RNAs have an exposed 3’-end. Substrate III contains a ^32^P-labeled RNA (*P-2) annealed to the template at a position where an additional RNA (P-3) is annealed downstream and blocks access to the 3’-end of the labeled RNA. Substrate III permits assessment of ExoN cleavage at a nick, as produced by nsp15 endonucleolytic cleavage. (**B**) ExoN does not initiate hydrolysis at a nick. The 10-nt and 15-nt RNAs are indicated. Reactions contained 0.5 µM ExoN (1:1) and 0.1 µM RNA and were quenched at the indicated times. The ^32^P-labeled RNAs (*P-3 and *P-2) in substrates I and II that have an exposed 3’-end were efficiently cleaved by ExoN. The ^32^P-labeled RNA (*P-2) in substrate III was not cleaved. (**C-E**) ExoN does not hydrolyze termini containing a 3’-phosphate or 2’,3’-cyclic phosphate. The structures of these modifications are shown in panel C. Schematic of dsRNA substrates containing 3’-phosphate and 2’,3’-cyclic phosphate modifications used are shown in panel D. Reactions contained 0.5 µM ExoN (1:1) and 0.1 µM RNA, were incubated for the indicated time, then quenched. Products are shown in panel E. The unmodified RNA is completely degraded; however, the 3’-phosphate and 2’,3’-cyclic modifications block excision by ExoN. Note, the 3’-phosphate and 2’,3’-cyclic modifications alter the apparent mobility of the RNA as it is more negatively charged and runs faster on the gel.

Cleavage by nsp15 leaves a 3’-end with a 2’-3’-cyclic phosphate (cycP) that can be hydrolyzed to a 3’-phoshphate (3’-PO4) and a 5’-end with a hydroxyl (**Fig. 2C**). Using approaches developed by the Shuman laboratory (**Fig. S4**) (33), we prepared RNAs to test as substrates for ExoN that contained either a cycP or a 3’-PO4 at the recessed end of the dsRNA (**Fig. 2D**). Neither of these RNAs served as substrates (**Fig. 2E**). So, even if the RNA downstream of the nick produced by nsp15 cleavage were removed, this product would be incapable of being degraded by ExoN without production of a 3’-hydroxyl.

### Nucleotide hydrolysis by ExoN occurs in a distributive manner, as expected for a proofreading exonuclease

All of the studies of nucleotide hydrolysis by ExoN published to date note the processive nature of the enzyme (18,23,28,44-51). In this case, processive means hydrolysis of multiples of ten nucleotides per binding event and is in contrast to a distributive enzyme, which would hydrolyze one or two nucleotides per binding event. Proofreading exonucleases associated with DNA polymerases have evolved to function in a distributive manner. This circumstance eliminates the mismatch without the need for re-synthesis of stretches of nucleic acid that were correctly basepaired, as would be the case if the exonuclease acted processively (26,27).

The ability to split the hydrolyzed strand into multiple components (**Fig. 2**), offered the opportunity to create a substrate with sufficient dsRNA to form a processive, elongation complex using the SARS-CoV-2 RTC (29,56,57) but at the same time permit hydrolysis by ExoN to be monitored with single-nucleotide resolution. The substrate used is shown in **Fig. 3A**. We evaluated utilization of this substrate by ExoN under two conditions. We refer to the first condition as substrate excess, which should reveal the length(s) of product formed by ExoN in a single binding event (left panel of **Fig. 3B**). We refer to the second condition as enzyme excess (right panel of **Fig. 3B**). Under these conditions and upon dissociation, the product of one enzyme in the reaction immediately serves as a substrate for the same or a second enzyme in a reiterative manner until the product dissociates into two single-stranded molecules. This latter condition is the most prevalent in the literature (23,44-47,50,58).

**Figure 3.**
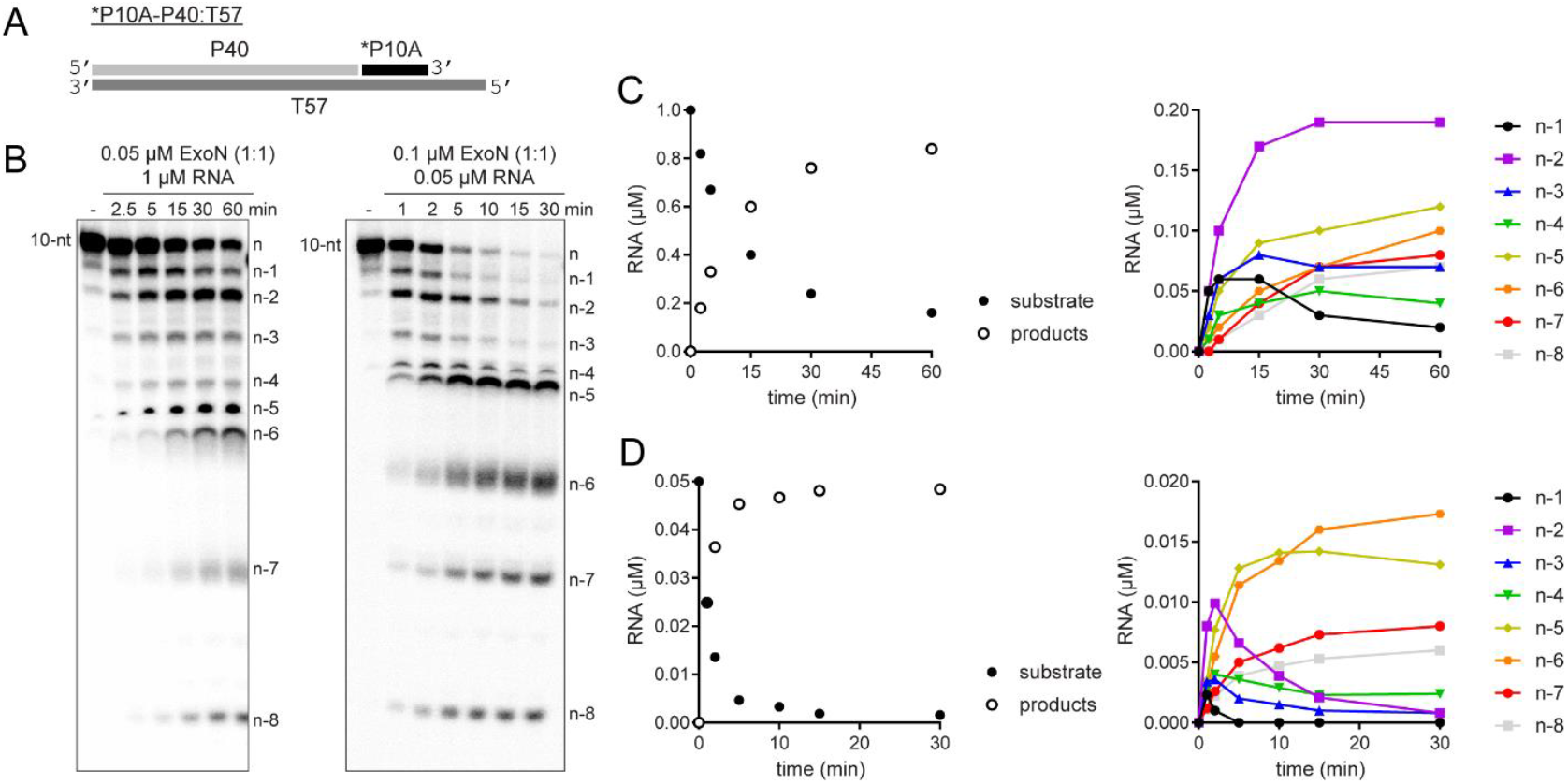
Monitoring ExoN-catalyzed RNA hydrolysis at single-nucleotide resolution. (**A**) Schematic of dsRNA substrate used to monitor ExoN activity at single-nucleotide resolution. (**B**) Reactions contained 0.05 µM ExoN (1:1) and 1 µM RNA or 0.1 µM ExoN (1:1) and 0.05 µM RNA, were incubated for the indicated time, then quenched. Products were resolved using denaturing 20 or 23% polyacrylamide gels. The RNA cleavage products, n-1 to n-8, are indicated. (**C**,**D**) Quantitative analysis of kinetics of substrate RNA utilization and/or product RNA formation. Formation of individual products is shown on the right. Panel C reports on the experiment in which RNA was in excess of enzyme; panel D reports on the experiment in which enzyme was in excess of RNA.

Under both conditions, the earliest products were primarily n-1 and n-2 in length (**Fig. 3B**). Quantitative analysis of both conditions showed consumption of more than 80% of the substrate during each reaction (left panels of **Figs. 3C** and **3D**). More than 50% of the substrate was consumed by the first time point when enzyme was present in excess (left panel of **Fig. 3D**), thus precluding an unambiguous assessment of precursor-product relationships (right panel of **Fig. 3D**). However, under the condition of substrate-excess, the kinetics suggested production of n-1 and n-2 at the same rate, with n-2 accumulating over the time course (right panel of **Fig. 3C**). Therefore, products shorter than n-3 likely arise from utilization of the n-1 and/or n-2 products.

### Existence of an equilibrium between nsp14 and nsp10 proteins

Structural studies of ExoN demonstrate unambiguously that the nsp14:nsp10 stoichiometry is 1:1 (19,22,28,45,46,48). However, there has been an assumption that the affinity of nsp14 for nsp10 is sufficiently high that complex never dissociates once formed. This interpretation is based on the fact that most published studies emphasize the nsp14:nsp10 stoichiometry rather than the concentration of each component (20,23,28,44-51,58). Because most published studies have used conditions of enzyme excess, any dependence of the reaction rate on nsp10 concentration would be masked without evaluating product formation on the msec-sec timescale (**Fig. S5**).

We evaluated the reaction kinetics, processivity, and substrate specificity of nsp14 as a function of nsp10 concentration (**Fig. 4**). The reaction rate increased as a function of nsp10 concentration, without apparent saturation at a concentration of 2 µM, and nsp14:nsp10 ratio of 1:20 (**Fig. 4A**). The primed-template-like dsRNA substrate was still favored over the ssRNA substrate (compare the 10-nt band in **Fig. 4A** to that in **Fig. 4B**). Finally, the distributive nature of the enzyme remained the same, with n-1 and n-2 products accumulating at early times for both substrates (**Fig. 4**).

**Figure 4.**
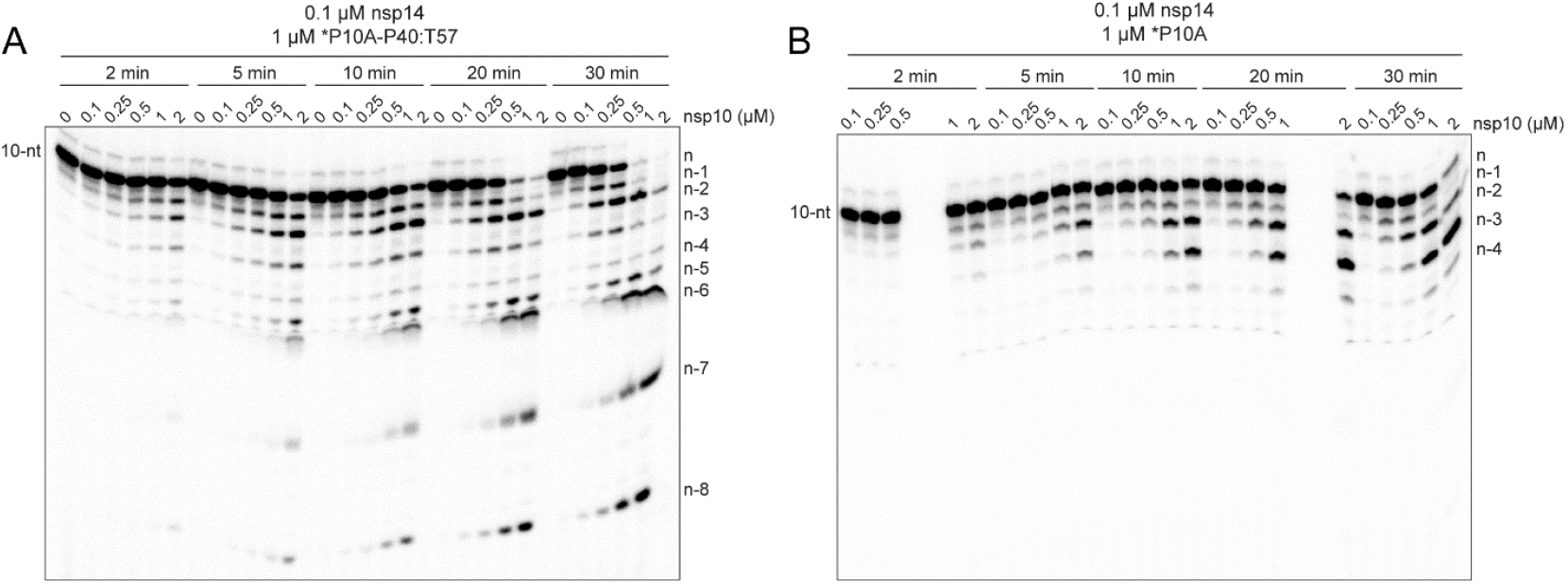
nsp10 stimulates hydrolysis without affecting processivity or substrate specificity. (**A**) Effect of nsp10 concentration on the kinetics of nsp14-catalyzed dsRNA hydrolysis. Reactions contained fixed nsp14 (0.1 µM) and varied nsp10 (0 – 2 µM) concentrations. In all cases, nsp14 was pre-mixed with nsp10 on ice 5 min prior to adding to the reaction. Hydrolysis of dsRNA substrate (*P10A-P40:T57) was monitored over a 30 min time course as indicated. Product analysis is shown. (**B**) Effect of nsp10 concentration on the kinetics of nsp14-catalyzed hydrolysis of ssRNA. Reactions were as in panel A except a ssRNA substrate (*P10A) was used. Product analysis is shown. Products shorter than n-4 are not observed when the 10-nt ssRNA is used.

An understanding of the kinetics and thermodynamics of the binding reaction between nsp14 and nsp10 leading formation of the nsp14-nsp10 complex is warranted. Such studies will require approaches to assess complex formation directly, rather than indirectly by monitoring exoribonuclease activity.

### The presence of the phosphorothioate Rp diastereomer at the scissile phosphodiester bond blocks hydrolysis by ExoN

Polymerases prefer use of the Sp diastereomer when a phosphorothioate is placed at the alpha position of a nucleoside triphosphate (59-63). Incorporation yields the Rp diastereomer at the resulting phosphorothioate bond (**Fig. 5A**), and the presence of this diastereomer can block hydrolysis by some exonucleases (64-68). We substituted the ultimate (PS110A in **Fig. 5B**), penultimate (PS210A in **Fig. 5B**), and antepenultimate (PS310A in **Fig. 5B**) phosphodiester bond with a phosphorothioate. Both diastereomers were present at a 50:50 ratio.

**Figure 5.**
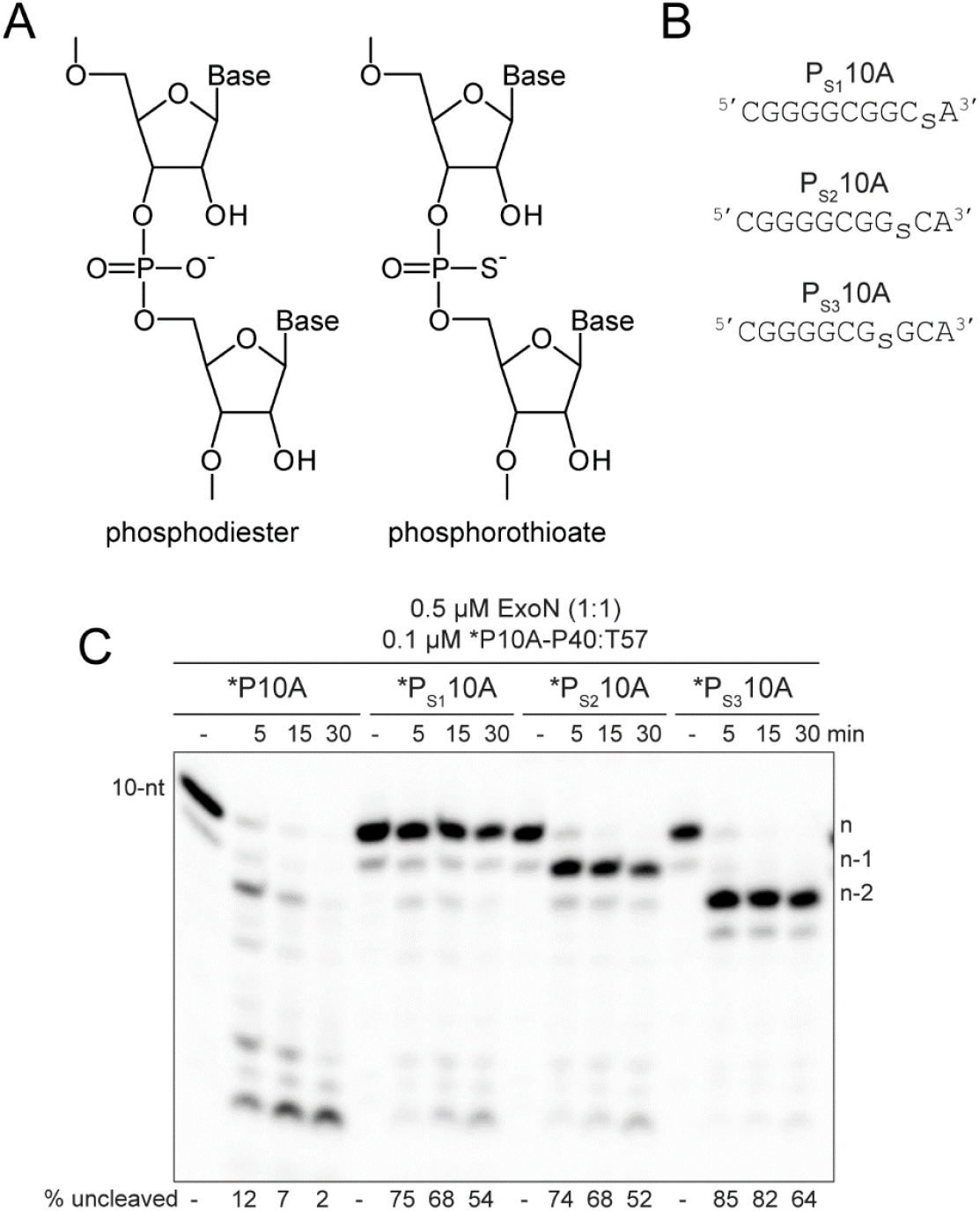
RNA hydrolysis by ExoN is inhibited by a phosphorothioate bond. (**A**) Structure of phosphorothioate. One of the non-bridging oxygens of the phosphodiester bond is replaced with a sulfur atom, creating Rp and Sp diastereomers. A 50:50 ratio of diastereomers were used in these experiments. (**B**) Schematic of phosphorothioate-substituted ssRNAs used. (**C**) Effect of phosphorothioate ExoN-catalyzed hydrolysis of dsRNA. Reactions contained 0.5 µM ExoN (1:1) and 0.1 µM of the indicated RNA, were incubated at the indicated times, then quenched. Product analysis is shown. Fraction of RNA remaining is indicated (% uncleaved). The unmodified RNA is completely degraded; however, ExoN is unable to hydrolyze beyond the phosphorothioate substitution, for half the RNA molecules at least. This observation is consistent with only a single diastereomer being inhibitory.

We assembled each phosphorothioate-substituted RNA into the *P10A-P40:T57 substrate to determine the impact of the phosphorothioate substitution on hydrolysis by ExoN compared to the control RNA (**Fig. 5C**). Under the conditions of enzyme excess, essentially all of the control substrate was consumed (*P10A in **Fig. 5C**). However, only 50% or so of the substrate was consumed when a phosphorothioate was present in the RNA (*PS110A, *PS210A, and *PS310A in **Fig. 5B**), suggesting that one diastereomer inhibits hydrolysis. Importantly, the size of the terminal product in the reaction was consistent with the position of the phosphorothioate: n, for *PS110A; n-1 for *PS210A; and n-2, for *PS310A (**Fig. 5B**).

To determine if the phosphorothioate Rp diastereomer was the inhibitory species, we used polymerase incorporation of nucleoside-5’-O-(1-thiotriphosphates) to produce RNA with only the Rp diastereomer at the scissile bond (**Fig. 6A**). For these experiments, we used the cryptic RdRp activity present in the mitochondrial DNA-dependent RNA polymerase (DdRp), POLRMT. For these experiments, we used the DdRp to produce an RNA product with the terminal scissile bond containing the phosphorothioate Rp diastereomer (**Fig. 6B**). After approximately 50% of the primer was extended to product, we added ExoN and monitored the fate of the product RNA by denaturing PAGE and phosphorimaging (**Fig. 6B**). Extension of only half of the product provides an internal control for the presence of ExoN, because the primer lacks a phosphorothioate bond. We used two different substrates. For both, the presence of the phosphorothioate Rp isomer completely blocked hydrolysis by ExoN (right panels in **Figs. 6C,D**), relative to both the unextended primer (internal control) and the comparable RNA product lacking the phosphorothioate (left panels in **Figs. 6C,D**). Quantitation is provided in **Figs. 6E,F**. Addition of the phosphorothioate to the α-position of an antiviral nucleotide analog should eliminate the possibility for excision by ExoN.

**Figure 6.**
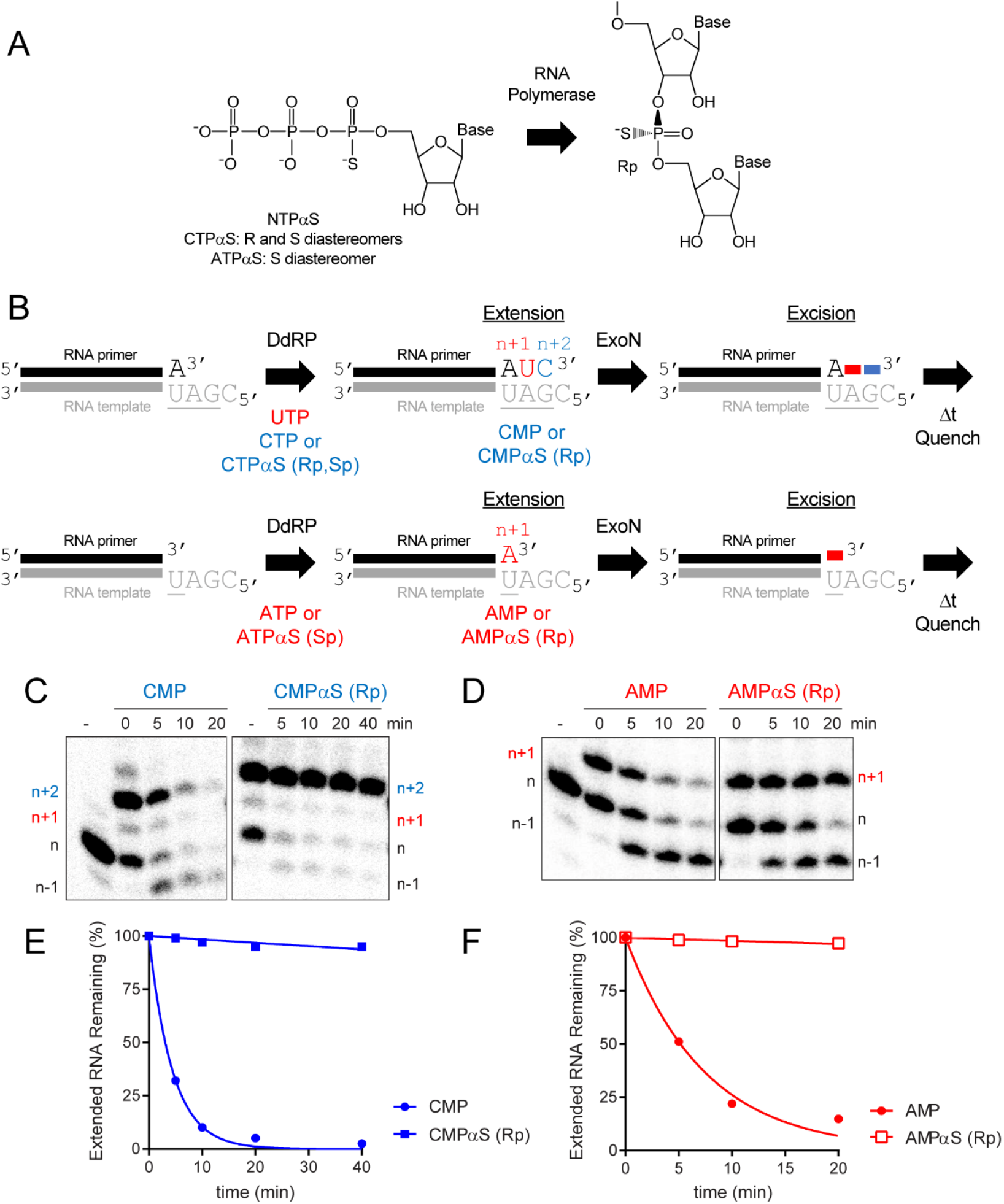
ExoN is inhibited by the Rp diastereomer at the phosphorothioate-substituted scissile bond. **(A)** The Sp diastereomer of a nucleoside-5’-O-thiotriphosphate is preferentially incorporated by RNA polymerases. Nucleophilic attack at the alpha phosphorous atom leads to inversion of configuration and creation of Rp diastereomer at the phosphorothioate bond. (**B**) Schematic of assay. Primer extension is initiated by adding an RNA polymerase in the presence of either UTP and CTPαS or ATPαS. CTPαS is a mixture of both the Rp and Sp diastereomers. ATPαS was obtained as the pure Sp diastereomer. Incorporation yields extended products. The last nucleotide to be incorporated is either CMP(αS) or AMP(αS). Once 50-75% of the primers were extended to the end, ExoN was added to the reaction. The reaction was monitored over time for hydrolysis. Unextended primer in reactions served as a useful control to demonstrate the presence of active ExoN in the reaction. (**C, D**) Analysis of reaction products by denaturing PAGE. Incorporation of CMP and AMP results in removal of the incorporated nucleotide by ExoN. Incorporation of CMPαS and AMPαS (Rp diastereomers) results in a terminated primer that cannot be cleaved by ExoN. (**E**,**F**) Kinetics of excision of CMP, CMPαS (Rp diastereomer) and AMP and AMPαS (Rp diastereomer) by ExoN. Data were fit to a single exponential. Rates are provided in **Table 2**.

### Composition of the terminal basepair modulates the kinetics of hydrolysis by ExoN

For proofreading exonucleases of DNA polymerases, it is not that the exonuclease exhibits a preference for a mispaired end relative to a properly paired end (26,27). Rather, a mispaired end is unstable in the polymerase active site, providing the opportunity for the mispaired end to partition to the exonuclease active site (26,27). To assess the impact of the terminal basepair on the kinetics of excision by ExoN, we designed two additional RNA substrates for ExoN. *P10G creates a terminal G:U mispair; *P10R creates a terminal remdesivir:U pair, which is equivalent to an A:U pair at the level of the hydrogen bonding potential between the bases (**Fig. 7A**). We evaluated the kinetics of hydrolysis of the various RNA substrates by ExoN under conditions in which the RNA substrate was present in excess of enzyme. Reaction products were resolved by denaturing PAGE and visualized by phosphorimaging (**Fig. 7B**). The observed rates of consumption of the RNA substrate and formation of all products were the same (**Fig. 7C**). Any difference that would be observed, however, would be related to hydrolysis of 3’-terminal nucleotide to produce the n-1 product and any subsequent excision reactions occurring prior to dissociation of ExoN. We performed a comprehensive analysis of the kinetics of formation of each product for each substrate (**Fig. 7D**). We observed a clear difference in the magnitude of the n-1 product that accumulated for each substrate (**Fig. 7D**). Such an observation would suggest that the rate of the step governing hydrolysis and/or rate of dissociation of the excised nucleotide product differ between the substrates. Interestingly, excision of remdesivir was substantially slower than the natural nucleotides as the n-1 product accumulated to the greatest extent for the P10R substrate (**Fig. 7D**).

**Figure 7.**
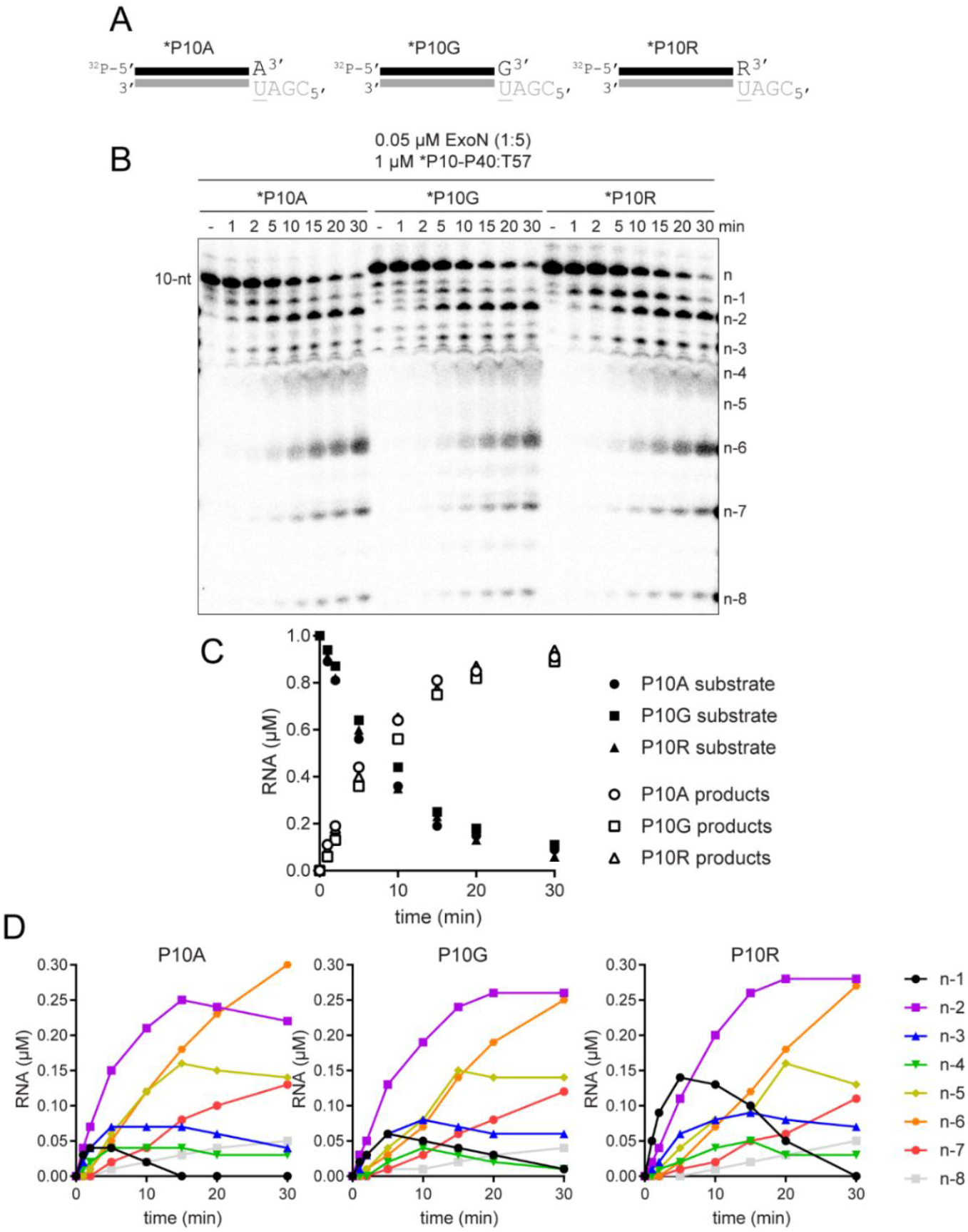
Terminal basepair-dependent differences in the kinetics of hydrolysis by ExoN. (**A**) Schematic of dsRNA substrates used to evaluate the influence of the terminal basepair on kinetics of excision by ExoN. The termini have a A:U, G:U or R:U basepair, where R represents remdesivir. (**B**) Effect of terminal basepair on hydrolysis by ExoN. Reactions contained 0.05 µM ExoN (1:5) and 1 µM of the indicated dsRNA substrate, were incubated at 30 °C for the indicated times, then quenched. Product analysis is shown. (**C**) Analysis of the kinetics of substrate utilization and product formation. Here, depletion of substrate and total product formation are monitored, showing no difference in the kinetics. (**D**) Analysis of the kinetics of formation of each individual product. Cleavage products (n-1 to n-8) were plotted as a function of time for each RNA substrate. The kinetics of formation and utilization of the n-1 product vary between substrates, suggesting that some, but not all steps of the mechanism are agnostic to the sequence and/or complementarity of the terminal basepair.

### Some modifications to the 3’-terminal ribose antagonize ExoN-catalyzed excision

To further our understanding of the structure-activity relationships of the 3’-terminal nucleotide, we prepared substrates in which the 2’- or 3’-hydroxyl was changed (**Fig. 8A**). Our standard substrate RNA is P10A. Unfortunately, not all modifications were available as an adenosine phosphoramidite. Therefore, some modifications were analyzed in the context of P10U (**Fig. 8A**). We monitored the kinetics of ExoN cleavage of modified RNAs. Products were resolved by denaturing PAGE and visualized by phosphorimaging (**Figs. S6** and **S7**). We plotted the concentration of total RNA products as a function of time for the two conditions used: excess substrate (**Fig. 8B**) or excess enzyme (**Fig. 8C**). The corresponding observed rates are presented in **Table 3**. The presence of a 3’-phosphate blocked hydrolysis (**Fig. 2E**). The presence of a 2’-fluoro had no effect. All other substitutions reduced the efficiency of cleavage, with the most effective being 3’-H, 2’-NH2 and 2’-H (**Figs. 8B**,**C** and **Table 3**). Together, these data suggest that it may be possible to identify substituents on the 3’-terminal ribose that would not impact utilization by the RTC but would inhibit excision by ExoN.

**Table 2.**
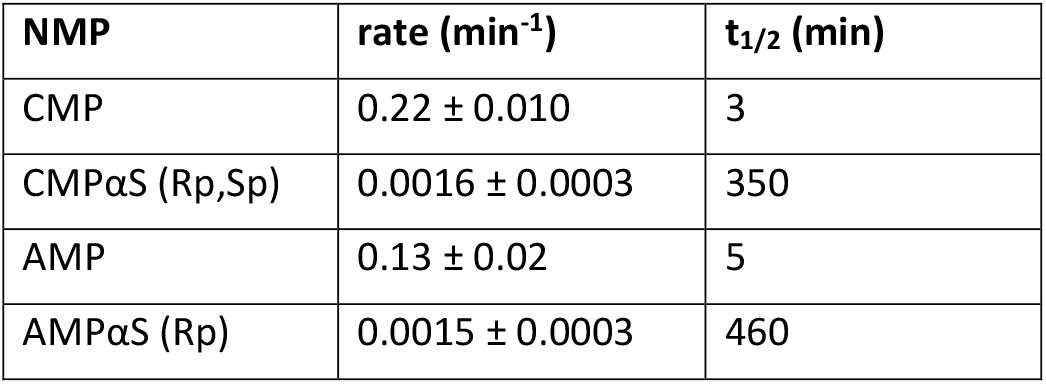
Kinetics of excision of the Rp diastereomer of CMPαS and AMPαS by ExoN. Data were fit to a single exponential to determine the observed rate of cleavage and half-life (t_1/2_).

| **NMP** | **rate (min<sup>-1</sup>)** | **$t_{1/2}$ (min)** |
| --- | --- | --- |
| CMP | $0.22 \pm 0.010$ | 3 |
| CMPαS (Rp,Sp) | $0.0016 \pm 0.0003$ | 350 |
| AMP | $0.13 \pm 0.02$ | 5 |
| AMPαS (Rp) | $0.0015 \pm 0.0003$ | 460 |

**Table 3.** Effect of modifications to the 3’-end on the kinetics of excision by ExoN. Data were fit either to a line (S>E) or to a single exponential (E>S) to determine observed rate of cleavage.

| modification | rate (min<sup>-1</sup>) | rate (min<sup>-1</sup>) |
| --- | --- | --- |
|  | S > E | E > S |
| unmodified | 0.87 ± 0.1 | 2.2 ± 0.2 |
| 2'-fluoro | 0.85 ± 0.3 | 1.7 ± 0.1 |
| 2'-O-methyl | 0.64 ± 0.02 | 0.46 ± 0.04 |
| 2'-deoxy | 0.41 ± 0.01 | 0.33 ± 0.02 |
| 2'-amino | 0.33 ± 0.01 | 0.15 ± 0.02 |
| 3'-deoxy | 0.035 ± 0.006 | 0.056 ± 0.008 |
| 3'-phosphate | no cleavage | no cleavage |

**Table 4.** Kinetics of excision of chain terminating cytidine analogs by ExoN. Data were fit to a single exponential to determine the observed rate of cleavage and half-life (t_1/2_).

|  | - SARS replicase<sup>a</sup> |  | + SARS replicase<sup>b</sup> |  |
| --- | --- | --- | --- | --- |
| **NTP** | **rate (min<sup>-1</sup>)** | **$t_{1/2}$ (min)** | **rate (min<sup>-1</sup>)** | **$t_{1/2}$ (min)** |
| CTP | 0.23 ± 0.01 | 3 | 0.0027 ± 0.0004 | 260 |
| UTP | - | - | 0.0046 ± 0.0002 | 150 |
| 3'-dCTP | 0.014 ± 0.002 | 50 | 0.0032 ± 0.0004 | 220 |
| ddhCTP | 0.089 ± 0.01 | 8 | 0.0018 ± 0.0002 | 390 |
| 2'-C-Me-CTP | 0.15 ± 0.02 | 5 | 0.0028 ± 0.0001 | 250 |

<sup>a</sup>Experiment shown in **Fig. 9**.
<sup>b</sup>Experiment shown in **Fig. 10**.

**Figure 8.**
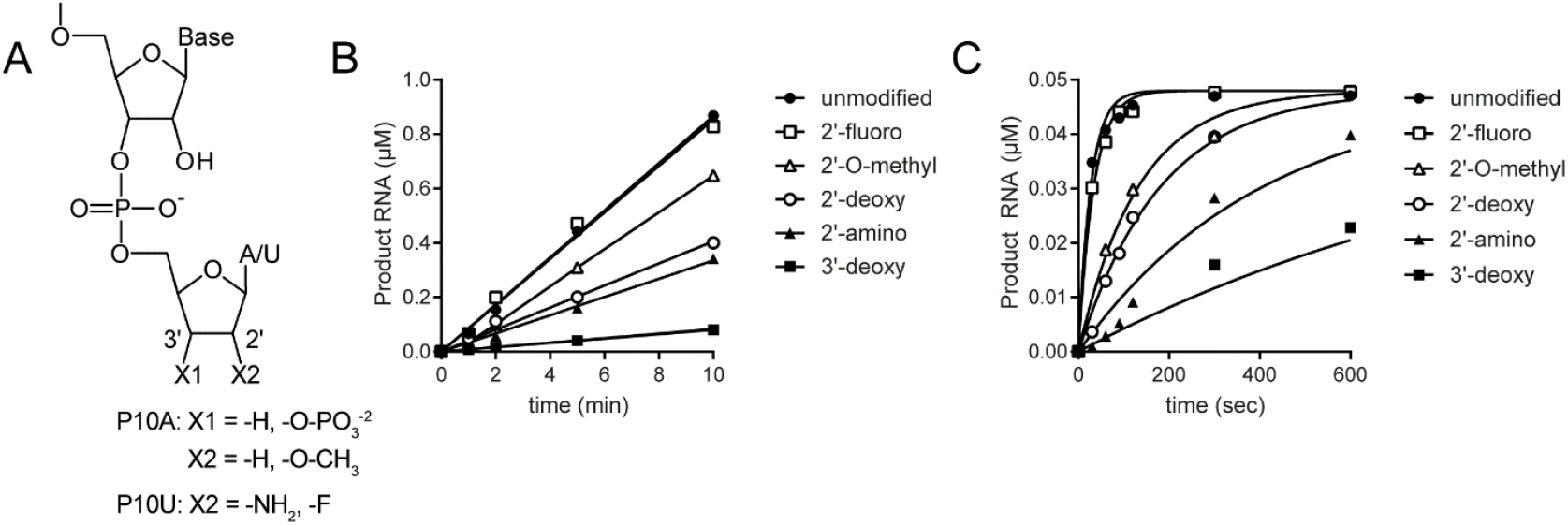
Structure-activity relationships for the ribose of the 3’-terminal nucleotide. (**A**) Modifications to the ribose of the terminal nucleotide studied here in the context of P10A: 3’-deoxy (-H), 3’-phosphate (-O-PO3), 2’-deoxy (-H), and 2’-O-methyl (-O-CH3); or P10U: 2’-amino (-NH2) and 2’-fluoro (-F). (**B**,**C**) Kinetics of cleavage of modified dsRNAs. In panel B, substrate was present in excess of enzyme. In panel C, enzyme was present in excess of substrate. The data were fit to a line (panel B) or to a single exponential (panel C). Reactions contained 0.1 µM ExoN (1:20) and either 1 or 0.05 µM dsRNA substrate and were quenched at the indicated times. The order in which the modified RNAs were cleaved is as follows: unmodified=2’-F > 2’-O-Me > 2’-deoxy > 2’-amino > 3’-deoxy > 3’-phosphate. The kinetics of cleavage using the 3’-phosphate RNA is not shown, because cleavage was not detected. The observed rate of cleavage of modified RNAs and fold difference from the unmodified RNA are shown in **Table 3**.

### ExoN efficiently excises the chain-terminating antiviral ribonucleotide produced by viperin, ddhC

The literature is rife with examples of the inability of obligate and non-obligate chain terminators to interfere with SARS-CoV-2 multiplication in cell culture (8,13,69-71), even the antiviral ribonucleotide, ddhCTP, produced by the antiviral protein, viperin (72). However, a 3’-dAMP terminated RNA is not a good substrate for ExoN (**Fig. 8**), and others have made similar observations for other 3’-dNMPs (28,73).

Given this apparent contradiction between the cell-based and biochemical experiments, we asked how ExoN would deal with the presence of the viperin product, ddhCTP, and the non-obligate chain terminator, 2’-C-Me-CTP (structures shown in **Fig. 9A**), when present at the 3’-end of RNA. Because synthetic RNA containing these analogs was not available, we made these RNAs biosynthetically by taking advantage of the cryptic RdRp activity present in the mitochondrial DNA-dependent RNA polymerase (DdRp, aka POLRMT, **Fig. 9B**). CTP and 3’-dCTP were used as controls to permit comparison to data obtained using synthetic RNA. After production of the CMP analog-terminated RNA, ExoN was added, and the reaction proceeded for the indicated time before quenching (**Fig. 9B**). Reaction products were resolved by denaturing PAGE and visualized by phosphorimaging (**Fig. 9C**) and quantified (**Fig. 9D**). The presence of the 2’-C-Me substituent had little to no effect on ExoN activity (2’-C-Me-CTP in **Figs. 9C,D**). In spite of the absence of a 3’-OH on ddhCTP, ddhCMP was excised from RNA almost as efficiently as CTP (ddhCTP in **Figs. 9C,D**). Together, the results presented in **Figs. 8** and **9** reveal complexity to the interaction between the 3’-terminal nucleotide and the ExoN active site that is not explained by simple docking experiments. A more in-depth characterization of the substrate specificity of ExoN in warranted.

**Figure 9.**
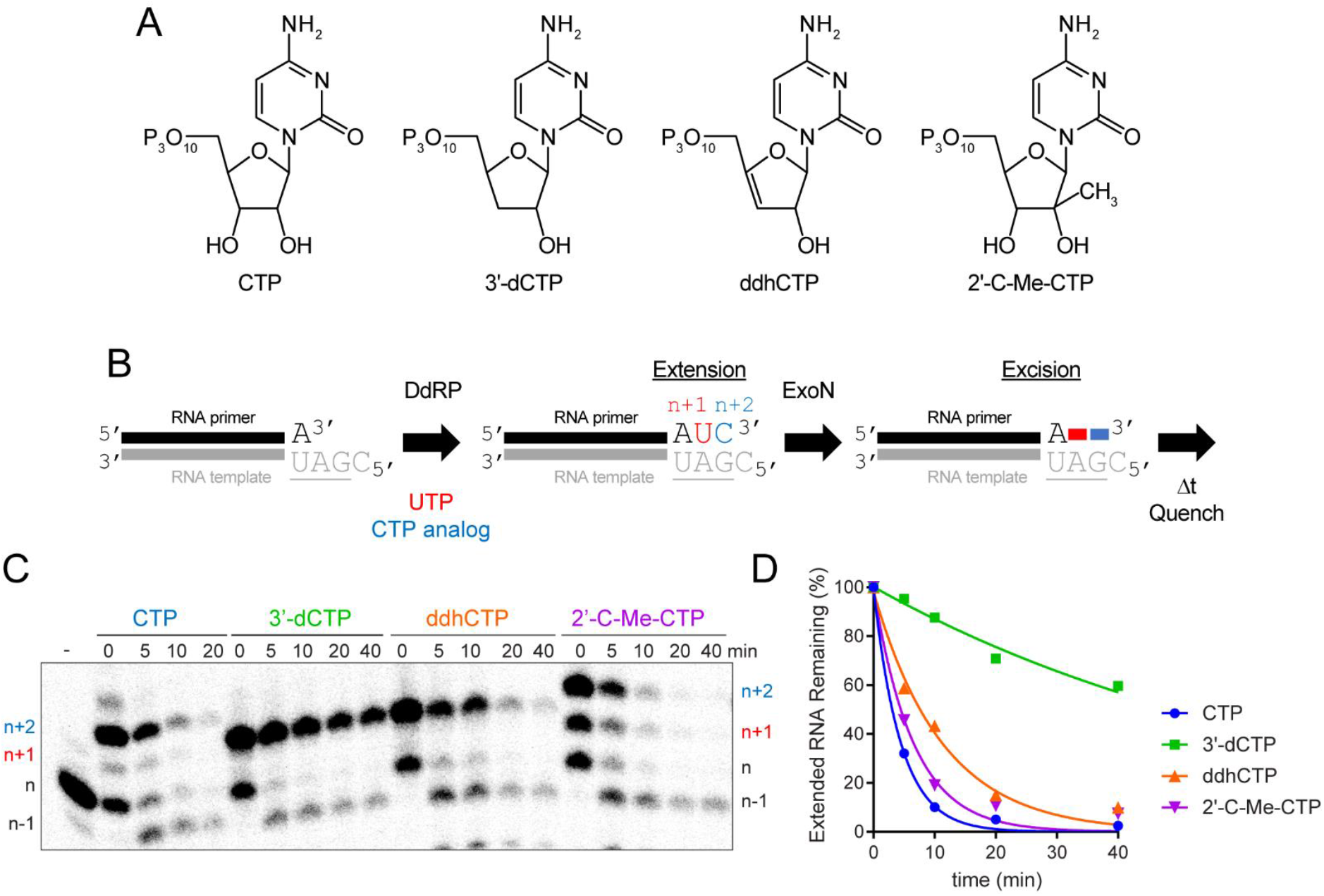
Viperin product (ddhCMP)-terminated RNA is a good substrate for ExoN but 3’-dCMP-terminated RNA is not. (**A**) Structure of nucleotide analogs used. (**B**) Schematic of the assay. Primer extension is initiated by adding an RNA polymerase in the presence of UTP and a CTP analog. Incorporation produces n+1 and n+2 products. The last nucleotide to be incorporated is the CTP analog. Once 50-75% of the primers were extended to n+2, ExoN was added to the reaction. The reaction was monitored over time for hydrolysis. Unextended primer in reactions served as a useful control to demonstrate the presence of active ExoN in the reaction. (**C**) Analysis of reaction products by denaturing PAGE. Only the 3’-dCMP-terminated RNA exhibited a delay in excision. (**D**) Kinetics of excision of CMP analogs. The quantitation revealed that only 3’-dCMP-terminated RNA exhibited a significant delay in the rate of excision relative to the CMP control, even though ddhCMP also lacks a 3’-OH. Data were fit to a single exponential. Rates are provided in **Table 4**.

### SARS-CoV-2 RTC prevents access of ExoN to the 3’-end nascent RNA

We have been able to assemble elongation complexes comprised of SARS-CoV-2 RTC and our split-primer:template substrate used for ExoN (**Fig. 10A**). Studies using similarly designed substrates with shorter duplexes fail to assemble a stable complex (data not shown). The ability to split the primer provides the advantage of being able to monitor extension at single-nucleotide resolution.

**Figure 10.**
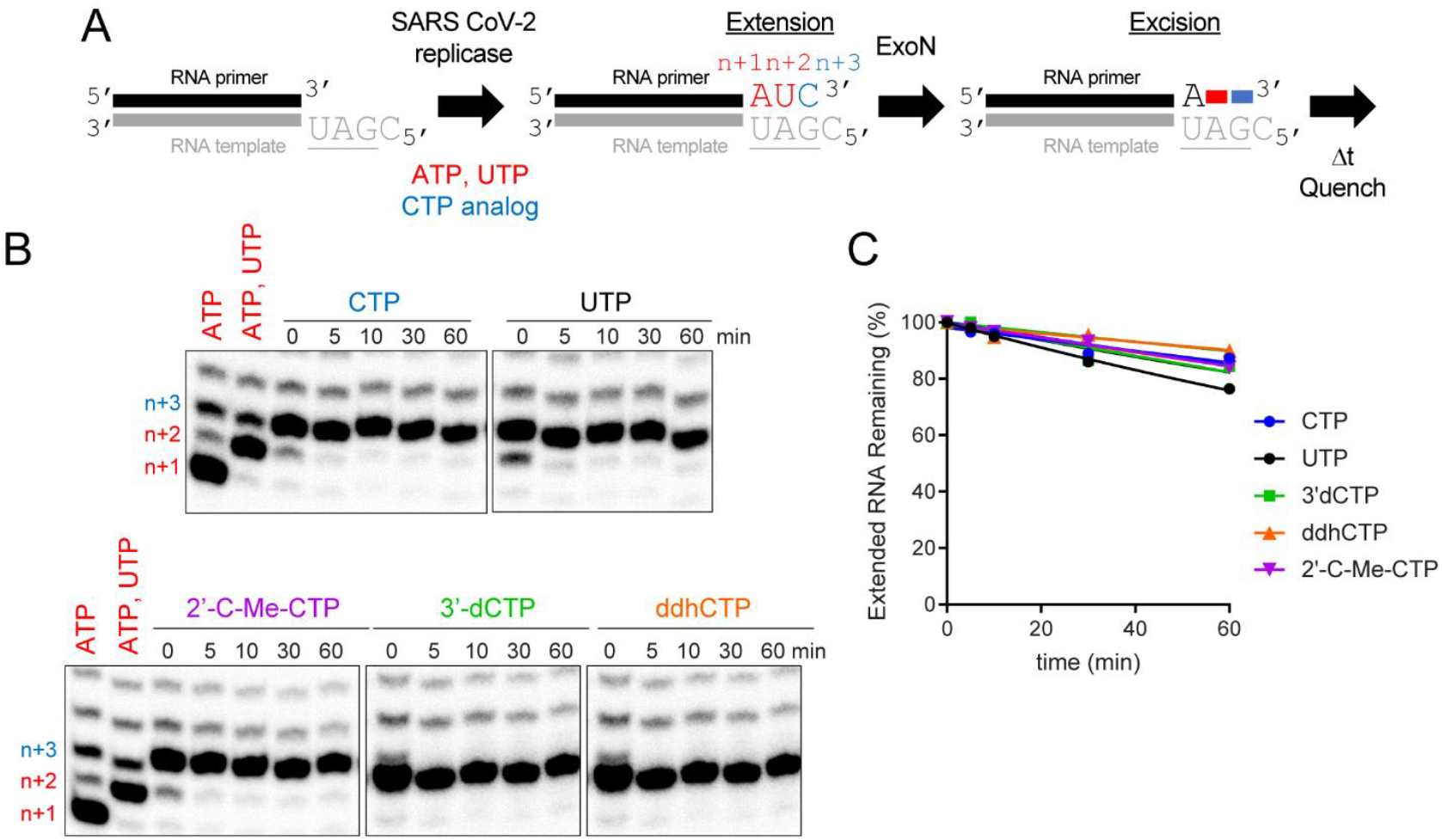
Stalled SARS-CoV-2 replication-transcription complexes block access of ExoN to RNA terminus. (**A**) Schematic of assay. Primer extension was initiated by adding the SARS-CoV-2 replicase-transcription complex (nsp7/nsp8/nsp12) in the presence of ATP and UTP or ATP, UTP, and CTP or a CTP analog. After an incubation time of 60 min, we added ExoN, incubated for the indicated time, and then quenched the reaction. (**B**) Effect of ExoN on stalled elongation complexes. Incorporation yields n+1, n+2, and n+3 products. When ATP, UTP, and CTP are present, the n+3 product has a properly basepaired 3’-end. The n+4 product is formed by misincorporation and has a mispaired 3’-end. When only ATP and UTP (low concentration, 1 µM UTP) are present, both the n+3 and n+4 products are formed by misincorporation and both products have mispaired 3’-ends. At high concentrations of UTP (100 µM), most of the products observed are at n+3 from misincorporation of U opposite G. ExoN was added to elongation complexes for the indicated time (5 – 60 min). Reaction products were visualized by phosphorimaging after denaturing PAGE. No change in the level of properly paired (n+3) or mispaired (n+4) elongation products was apparent for any of the reactions performed, consistent with the replicase remaining bound to both products and obstructing access by ExoN. (**C**) Quantitation of reaction products shown in panel B. The data were fit to a line. Rates are provided in **Table 4**.

Assembly and incubation of a complex with a correctly basepaired 3’-end for 60 min (**Fig. 10A**; 0 min **Fig. 10B**) remains stable to challenge with ExoN for at least an additional 60 min (**Fig. 10A**; CTP in **Figs. 10B,C**). Judicious selection of substrate nucleotides and/or analogs permitted us to exploit this system to create elongation complexes with different 3’-ends. This experimental design permitted us to determine if the nature of the 3’-terminal basepair under investigation influenced the stability of assembled SARS-CoV-2 RTC and/or accessibility of ExoN. Neither a mispaired end (UTP in **Fig. 10B,C**) nor an end containing a chain terminating analog (2’-C-Me-CTP, 3’-dCTP, ddhCTP in **Fig. 10B,C**) changed the sensitivity to ExoN, so by inference did not change stability with the SARS-CoV-2 RTC either.

Together, these data show that the SARS-CoV-2 RTC creates a physical block to cleavage and that the composition of the terminal basepair does not trigger a response by the SARS-CoV-2 RTC that facilitates access by ExoN.

## Discussion

Coronaviruses encode a 3’-5’ exoribonuclease (11,13-15). A complex of the nsp10 and nsp14 proteins at a stoichiometry of 1:1 forms the active enzyme (19,22,28,46,48). The nsp14 subunit harbors the catalytic residues; however, this subunit fails to achieve a stable conformation competent for RNA binding and catalysis in the absence of nsp10 (22). Therefore, we have referred to the active complex as ExoN instead of the nsp14 subunit alone. The biological function of ExoN is proofreading, correction of errors introduced by the viral polymerase (11-14). ExoN also contributes to excision of antiviral nucleotides, as the sensitivity of the virus to certain antiviral nucleotides increases in the absence of exoribonuclease activity (24). This latter activity complicates the use of some conventional antiviral nucleotides for the treatment of coronavirus infection. The goal of this study was to discover strategies to interfere with excision of nucleotides by ExoN and thereby inform the development of nucleotide analogs with greater efficacy against coronaviruses.

The catalytic cycle of ExoN includes binding of the RNA substrate to the enzyme followed by hydrolysis (**Fig. 11A**). There is now a consensus that the RNA substrate preferred by ExoN is one that would also be preferred by the replication-transcription complex (RTC), a primed-template (**Fig. 1**) (20,28,48). The number of nucleotides hydrolyzed per binding event was not known before this study. This gap existed for two reasons. First, the design of the RNA substrates used in these studies made difficult – if not impossible – the monitoring of product formation with single-nucleotide resolution (23,28,44,46-50,58). Second, ExoN was present in substantial excess over the RNA substrate, thus requiring monitoring of hydrolysis on the millisecond timescale to observe a product one nucleotide shorter than the substrate, which was never done (23,28,44,46-50,58). The inability of ExoN to hydrolyze nucleotides at a nick permitted us to distribute the overall requirement for dsRNA over multiple segments, which enabled product analysis at single-nucleotide resolution (**Figs. 2A,B**). While performing the experiments under conditions of enzyme excess may have some utility when evaluating inhibitors (44), only conditions of substrate excess can provide information on the number of nucleotides hydrolyzed per binding event when quenching reactions manually (**Fig. 11A**). ExoN excised only one or two nucleotides per binding event (**Fig. 3**). The distributive nature of ExoN makes sense, because a processive enzyme would then require the RTC to resynthesize RNA that lacked any damage.

**Figure 11.**
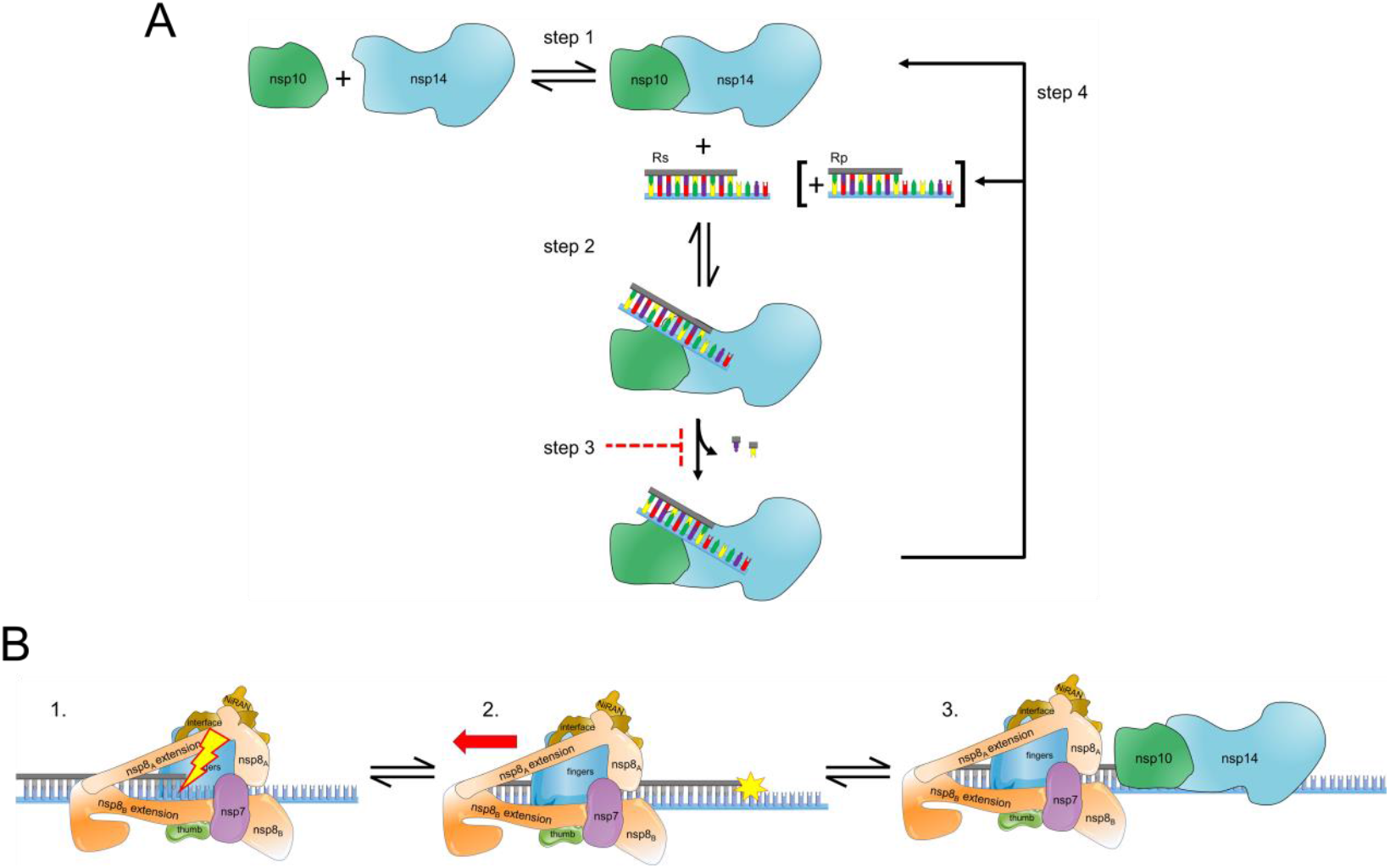
Hypothetical model for ExoN-catalyzed excision. (**A**) Excision uncoupled to RNA synthesis. The active exoribonuclease, referred to here as ExoN, is a complex of two subunits: nsp10 (regulatory) and nsp14 (catalytic). The concentration of ExoN therefore depends on the value of the dissociation constant for this binding reaction, as well as the concentrations of nsp10 and nsp14. Most studies to date have ignored this equilibrium. Under steady-state conditions in which the RNA substrate (RS) is present in excess of ExoN, RS binds, catalysis occurs, releasing one or two nucleotides, followed by dissociation of the RNA product (RP). Under these conditions, the rate-limiting step is likely release of RP. Modifications to the terminal nucleotide that reduce the rate of hydrolysis to values substantially lower than the rate of RP dissociation should interfere with excision. RP consumption depends on its ability to compete with RS, for example when more than 20% of the substrate is consumed or when ExoN is present in excess of substrate and more than a single turnover occurs. The possibility also exists for product dissociation to be driven by dissociation of the nsp10 and nsp14 subunits. (**B**) Excision coupled to RNA synthesis. Introduction of mispaired nucleotides or chain terminators by the polymerase complex does not lead to dissociation, blocking access of the 3’-end for repair by ExoN. The polymerase complex must therefore be actively dislodged from the terminus to access the 3’-end. Once ExoN binds, only one or two nucleotides will be removed.

For a nucleotide analog to resist excision, the rate constant for excision (Step 2 in **Fig. 11A**) needs to be slower than the rate constant for dissociation of ExoN from the 3’-terminus (Step 4 in **Fig. 11A**). RNA product dissociation is likely the rate-limiting step for this reaction under steady-state conditions (**Fig. 11A**). Both the 2’- and 3’-hydroxyls contribute to the efficiency of excision, as modifications to either appear to reduce the rate of catalysis to levels at or below the rate constant for dissociation (**Fig. 8, Table 3**). Our assumption is that when the rate of product formation is the same under conditions of excess substrate and excess enzyme, then the rate measured is catalysis (**Table 3**). The 3’-position was most important. Loss of the 3’-OH made catalysis rate limiting (**Fig. 8, Table 3**); addition of a 3’-phosphate eliminated turnover altogether (**Table 3**). The SARS-CoV-2 RTC has no problem utilizing nucleotide substrates lacking a 3’-OH (8,74). The issue with most analogs is the inability to compete with natural nucleotide pools (8,74). Remdesivir does not suffer this problem (8,74). Perhaps elimination of its 3’-OH will bolster its anti-coronavirus activity.

The ability of 3’-dCMP to antagonize ExoN-mediated hydrolysis so effectively was not expected, although several other groups have now reported this observation (28,73). Our previous studies showed that the antiviral nucleoside, ddhC, does not interfere with SARS-CoV-2 multiplication in cells, even though the RTC utilized the nucleotide, ddhCTP, quite readily (8). ddhC lacks a 3’-OH, but ExoN readily excised ddhCMP from the 3’-end of RNA (**Fig. 9**). We conclude that the requirement for a 3’-OH is not absolute but is context dependent. The context is likely associated with the ribose conformation. The ribose component of normal nucleotides are not planar and possess “sugar pucker” with the 3’-*endo* conformation being favored for ribonucleosides. The ribose of ddhC on the other hand is forced into a planar conformation because of the double bond between carbons 3’ and 4’ (**Fig. 9**). Structural studies will be required to provide an explanation for the context-dependent requirement for the 3’-OH.

In retrospect, it is not surprising that coronaviruses have evolved to evade the inhibitory activity of ddhCMP-terminated RNA. ddhCTP is induced by infection (72,75). Indeed, the presence of ddhCTP in cells presents a strong selective pressure for acquisition of an exoribonuclease, especially given the efficiency with which the RTC utilizes this antiviral nucleotide (8). The presence of an exoribonuclease may have also contributed to the polymerase retaining the ability to utilize ddhCTP. The evolutionarily related enteroviral polymerases have lost the ability to utilize ddhCTP (72). Some coronaviruses deal with loss of the exoribonuclease better than others(23), perhaps the efficiency with which the polymerases utilize ddhCTP has something to do with this. It would be interesting to determine the requirement of the exoribonuclease in cells lacking viperin, the enzyme that produces ddhCTP.

The most efficient strategy to interfere with nucleotide excision is to introduce a non-hydrolyzable scissile bond. Phosphorothioate (P-S) substitutions at the scissile phosphodiester bond inhibit excision by DNA exonucleases, usually requiring a specific stereoisomer (SP or RP) (64-68,76). Here we show that the presence of a phosphorothioate in the RP configuration of the scissile phosphodiester bond inhibits hydrolysis by ExoN (**Figs. 5** and **6** and **Table 2**). The ability to block cleavage of the terminal nucleotide rules out the existence of ExoN-associated endonuclease activity as suggest recently (44). Introduction of the P-S-substituted bond at ultimate, penultimate, and/or antepenultimate position will create substrates to facilitate elucidation of the details of ExoN-catalyzed hydrolysis by limiting the extent to which a product can be consumed. Current strategies to synthesize P-S-substituted oligonucleotides yield diastereomeric mixtures at phosphorous, but stereoisomer-specific solutions are on the horizon (76).

Another interesting possibility for use of the P-S substitution is the creation of non-hydrolyzable antiviral ribonucleotide analogs. Most clinically used monophosphorylated ribonucleotide analogs are readily delivered to cells as prodrugs that require intracellular activation by histidine triad nucleotide-binding protein 1 (HINT-1) (77,78), the so-called ProTide strategy (7,79-81). Similar strategies can be used for P-S-substituted ribonucleoside monophosphates. It is clear that HINT-1 can activate such compounds in cells, although the kinetics of activation are slower than conventional ProTides (77). HINT-1 is known to convert P-S-substituted NMPs to the natural NMP (77,78). The efficiency of this side reaction is likely tunable as the nature of the base and sugar contribute to the rate of sulfur removal (77). Side reactions such as these could also be alleviated by using P-S-substituted di- or triphosphorylated prodrugs (82). The pursuit of non-hydrolyzable ribonucleotide analogs is a guaranteed solution to the problem of excision of antiviral ribonucleotides by ExoN.

The majority of our studies focused on excision independent of RNA synthesis. An early report on ExoN from SARS-CoV suggested that co-transcriptional repair could occur in vitro (45,47). These studies used primed-templates of insufficient length to form a stable elongation complex (45,47). So, just as we were able to see “co-transcriptional” repair using POLRMT (e.g., **Fig. 9**), this repair required dissociation of POLRMT to permit ExoN access to the 3’-end. Formation of a stable elongation complex using the SARS CoV-2 RTC requires a duplex of 50-bp or greater (29,56,57). The ability to split the primer into 40-nt and 10-nt fragments permitted us to monitor extension by the RTC and excision by ExoN at single-nucleotide resolution in the same reaction (**Fig. 10**). Interestingly, assembling the SARS CoV-2 RTC on these primed templates and forcing the complex to make errors was insufficient to get the complex to dissociate and expose the 3’-end to ExoN (**Fig. 10**). ExoN removed these same termini in the presence of POLRMT (**Fig. 9**).

We conclude that the SARS-CoV-2 RTC alone does not sense the presence of a mispair, a non-natural nucleotide, or the pause caused by a chain terminator (panel 1 in **Fig. 11B**). A more active mechanism must exist to remove the RTC from the 3’-end, perhaps another viral factor pushes the RTC out of the way (panel 2 in **Fig. 11B**). The nsp13-encoded 5’-3’ RNA helicase may function in this capacity. nsp13 associates with the RTC (56,57). While incorporating nucleotides, the RTC may overcome helicase action. When stalled, however, the helicase is situated to push the RTC backwards (56,57). We suggest two potential scenarios: one where the replicase backtracks and one where the replicase is displaced backward. In the former, the product strand exits through the NTP channel, revealing the product strand 3’-end for mismatch correction by ExoN (cite Malone et al., PNAS 2021, Chen et al, Cell 2020, Liming et al., Cell 2021). In the latter, the displacement of the replicase leaves the duplex strand annealed and accessible to ExoN for mismatch correction (panel 3 in **Fig. 11B**). In both cases, establishing a tug of war between the polymerase and helicase in the RTC would favor exposure of any 3’-end that impedes forward motion of the polymerase. In such a scenario, access would regulate processing by ExoN and may explain the lack of a strict specificity for a mispaired end. So, what would happen in the presence of a catalytically inactive ExoN? It is likely that the complex would associate with the 3’-end, as illustrated in panel 3 of **Fig. 11B**, and create a physical block to synthesis by the RTC. It would be interesting to determine the extent to which substitutions that impair association of ExoN with the RNA duplex phenocopy those that preclude catalysis.

Remdesivir is one nucleotide analog that shows efficacy against SARS-CoV-2 in humans (1). The mechanism of action clearly relates to perturbed dynamics of the RTC (8,30,32,83). Whether or not some perturbations can occur immediately or only when present at the n+4 position has not been completely resolved. If remdesivir-terminated RNA were ever present, then our results would suggest that attempts at excision by ExoN would not be as facile as other nucleotide pairs (**Fig. 7**). This particular observation highlights the utility of monitoring hydrolysis by ExoN with single-nucleotide resolution.

The structure of ExoN reveals a complex of nsp10 and nsp14 at a 1:1 stoichiometry (19,22,28,45,46,48). However, such an observation does not infer picomolar affinity. We make this statement because the vast majority of the published studies on SARS-CoV-2 ExoN emphasize the stoichiometry of nsp10:nsp14 used without any consideration of the concentrations of each protein present. Our studies suggest that the dissociation constant for the nsp10-nsp14 complex is in the micromolar range (**Fig. 4**). Because our major conclusions derive from studies of the steady state at saturating concentrations of RNA, knowledge of the precise concentration of ExoN is not essential. However, conclusions from single-turnover experiments would be compromised. Direct studies of the nsp10-nsp14 binding equilibrium (Step 1 in **Fig. 11A**) will be essential to elucidation of the kinetic and chemical mechanisms.

Together, we describe a robust system for evaluating co-transcriptional proofreading of the SARS-CoV-2 polymerase and associated factor by the exoribonuclease. The framework established here will hopefully help the field at large by highlighting some of the key gaps in our understanding of the mechanism of ExoN-catalyzed nucleotide excision. Addressing these gaps will yield a clear picture of the structure-dynamics-function relationship of ExoN that can be used to guide design of ExoN-resistant nucleotide analogs.

## Data Availability

All data is available upon request.

## Acknowledgements

Our thanks to Roel Fleuren and Efra Rivera-Serrano for their contributions to the production of figures and models. We thank Stewart Shuman for providing the expression plasmid for RtcA.

