## Supplemental Data for "Interfering with nucleotide excision by the coronavirus 3’-to-5’ exoribonuclease"

### Supplementary Data

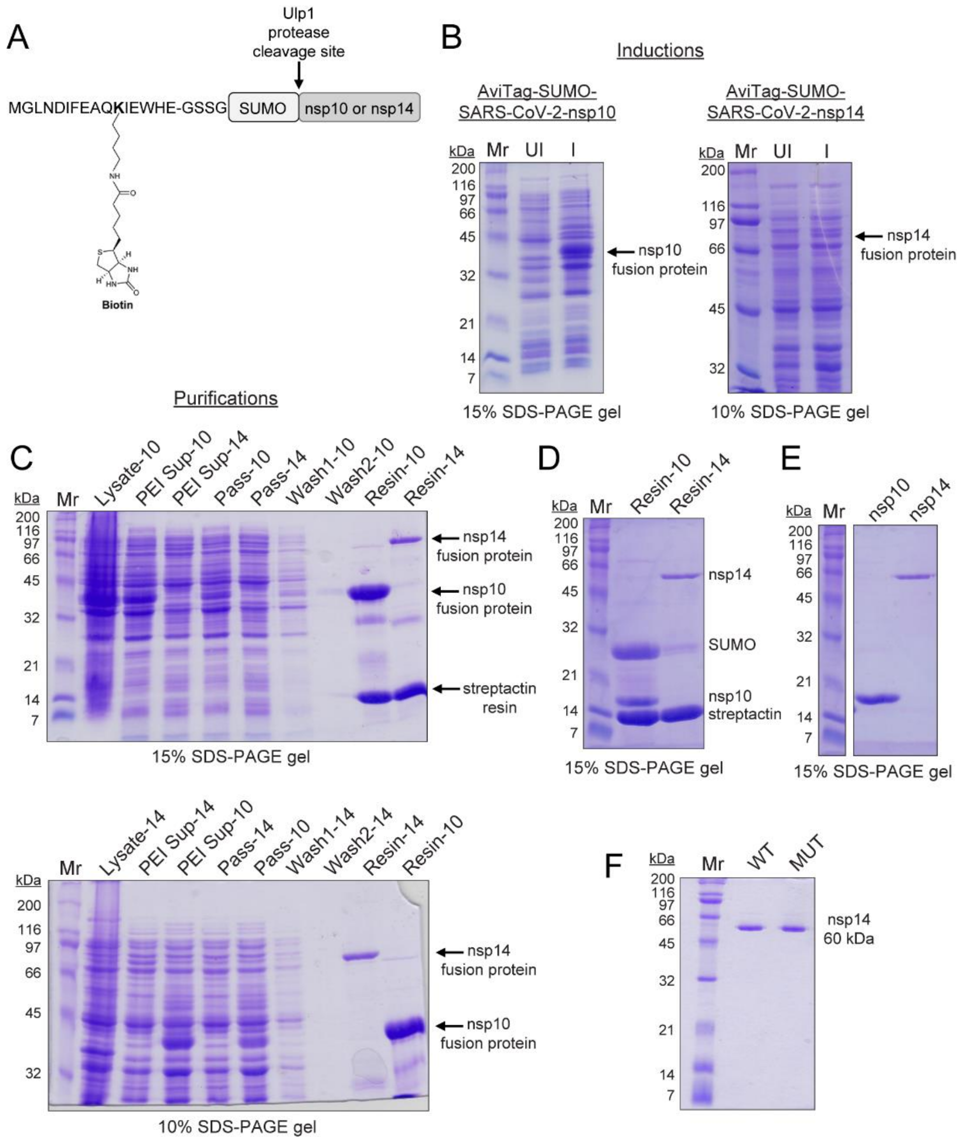


**Figure S1. Expression and purification of SARS-CoV-2 ExoN proteins: nsp10 and nsp14.** (**A**) Schematic of bacterial expression constructs used to express and purify SARS-CoV-2 nsp10 and nsp14. We employed AvitTag-SUMO fusion proteins for expression and purification of nsp10 and nsp14. The AviTag is a short 15 amino acid sequence (GLNDIFEAQKIEWHE) that can be specifically biotinylated on the lysine residue by the biotin ligase, BirA. The biotinylated AviTag allows affinity purification of the fusion protein to be isolated using streptavidin or streptactin resins. The SUMO fusion can be cleaved by the SUMO specific protease, Ulp1, which results in production of authentic, untagged target proteins. (**B**) SDS-PAGE analysis of bacterially expressed nsp10 and nsp14 SUMO fusion proteins. Shown are a 15% and 10% polyacrylamide gels with uninduced and induced samples for the Avitag-SUMO-SARS-CoV-2-nsp10 and Avitag-SUMO-SARS-CoV-2-nsp14 proteins. Proteins were expressed by IPTG at 25 $^{o}$C and 15 $^{o}$C as detailed in materials and methods. The position of the induced SUMO fusion proteins are indicated. Broad-range molecular weight markers (Mr) and corresponding molecular weights are indicated. Gel was stained with Coomassie. (**C**) SDS-PAGE analysis of samples from the purification of nsp10 and nsp14 SUMO fusion proteins are shown on 15% and 10% polyacrylamide gels with samples from each step of the purification. Samples loaded are: lysate, PEI supernatant, streptactin column pass, streptactin column washes, and streptactin resin with bound protein for nsp10 and nsp14 samples. The positions of the SUMO fusion proteins are indicated. The modified resin, streptactin, is also indicated. The resin was boiled in sample buffer prior to loading the gel. (**D**) SDS-PAGE analysis of samples of the streptactin resin with bound nsp10 and nsp14 SUMO fusion proteins after incubation with the SUMO protease, Ulp1. Both SUMO fusion proteins were completely cleaved by Ulp1. Cleavage results in the production of untagged full-length nsp10 and nsp14 proteins. The position of nsp10, nsp14, and cleaved SUMO protein are indicated. (**E**) SDS-PAGE analysis of nsp10 and nsp14 protein samples after Ulp1 cleavage and column wash to elute the untagged SARS-CoV-2 nsp10 and nsp14 proteins. The cleaved nsp10 and nsp14 proteins were effectively washed off the streptactin resin after cleavage resulting in isolation of the purified proteins. (**F**) Comparison of nsp14 WT and MUT (D90A E92A) purified proteins.

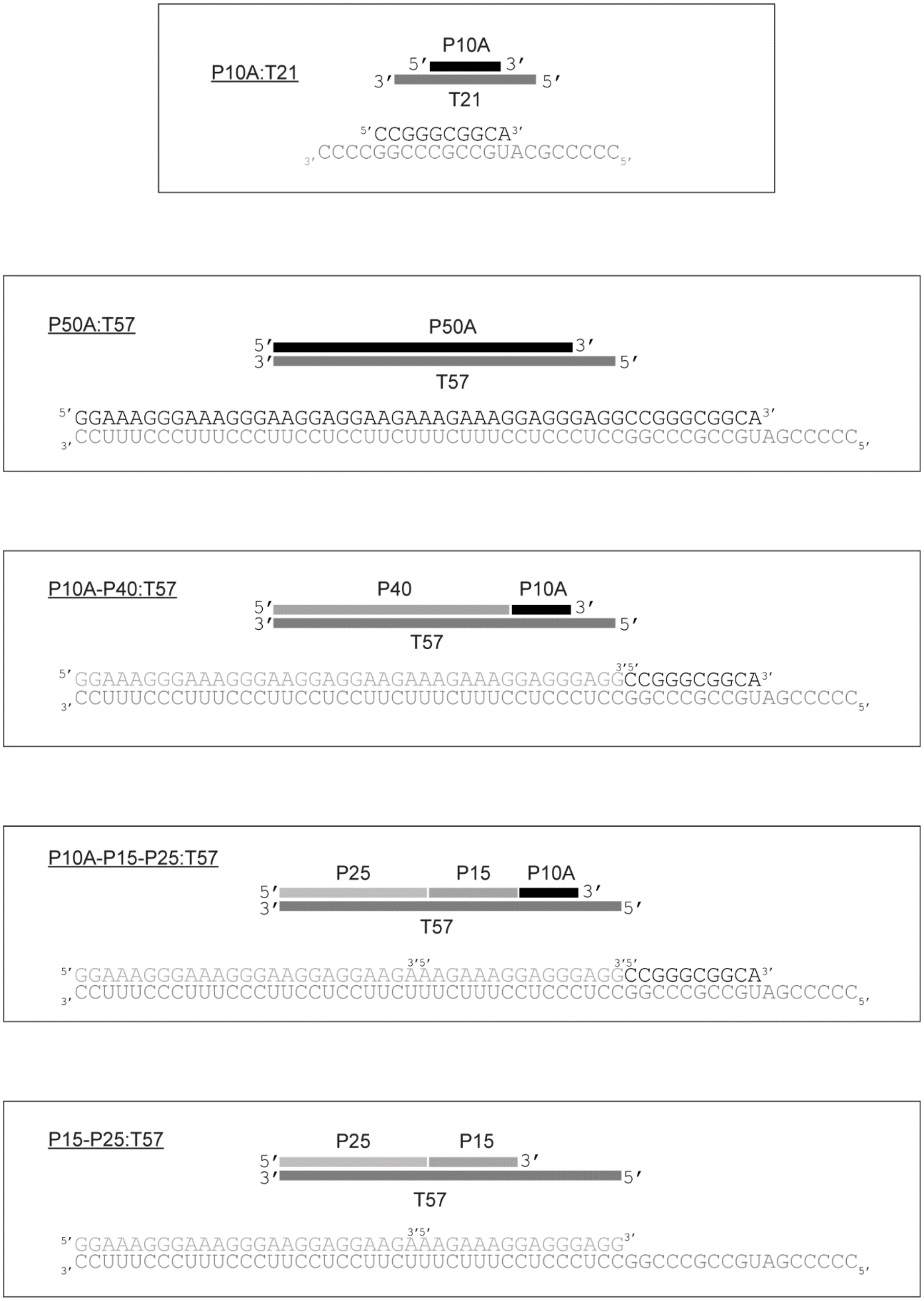


**Figure S2. Schematic of dsRNA substrates used in this study.** Shown are the dsRNA primed-template like substrates used in this study. RNA sequences are listed in Table 1.

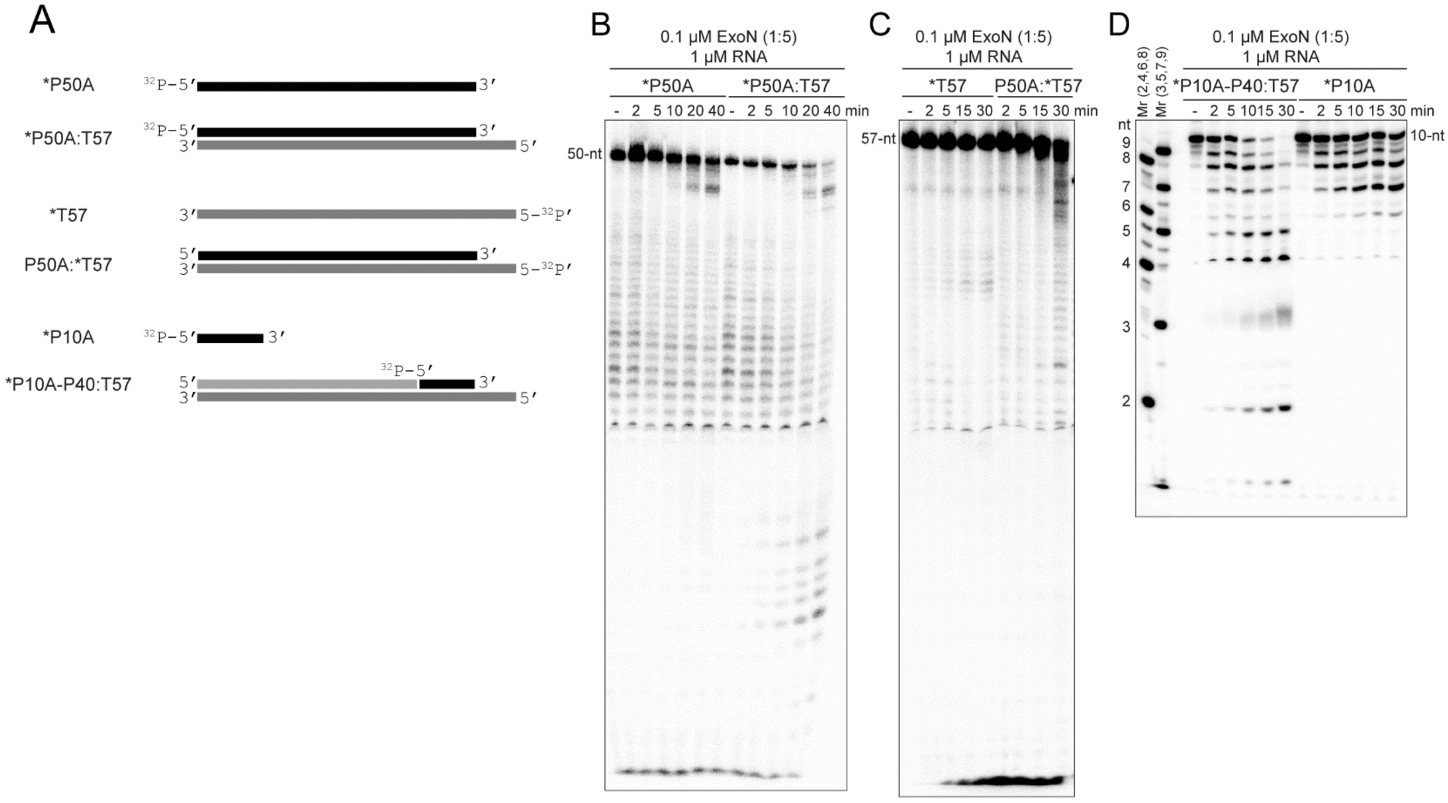


**Figure S3. ExoN cleaves primed-template-like substrates more efficiently than ssRNA, and a split $^{32}$P-labeled RNA primer on longer substrates allows for easier visualization of exonucleolytic cleavage products.** (**A**) Schematic of RNA substrates for evaluating SARS-CoV-2 exoribonuclease activity. RNA sequences are listed in Table 1 and RNA duplexes are shown in Figure S2. RNAs were $^{32}$P-end labeled, indicated by an asterisk (*), to monitor reaction products resolved by denaturing PAGE and visualized by phosphorimaging. (**C**) Analysis of reaction products by denaturing PAGE from using the RNAs depicted in panel A. The 50-nt, 57-nt and 10-nt $^{32}$P-labeled RNAs are indicated. Reactions contained 0.1 μM ExoN (1:5) and 1 μM RNA and were quenched at the indicated times. In both cases, the dsRNA primed-template like substrates were cleaved more efficiently than single-strand RNAs. The use of the 10-nt $^{32}$P-labeled RNA primer (*P10A) allows for easier visualization and better separation of the cleavage products compared to the 50-nt $^{32}$P-labeled RNA primer (*P50A). Note, the dsRNA substrates are essentially the same, however two RNAs (P10A-P40) are annealed to T57 instead of just one RNA (P50). $^{32}$P-labeled single strand RNA markers (2 to 9 nts) are indicated.

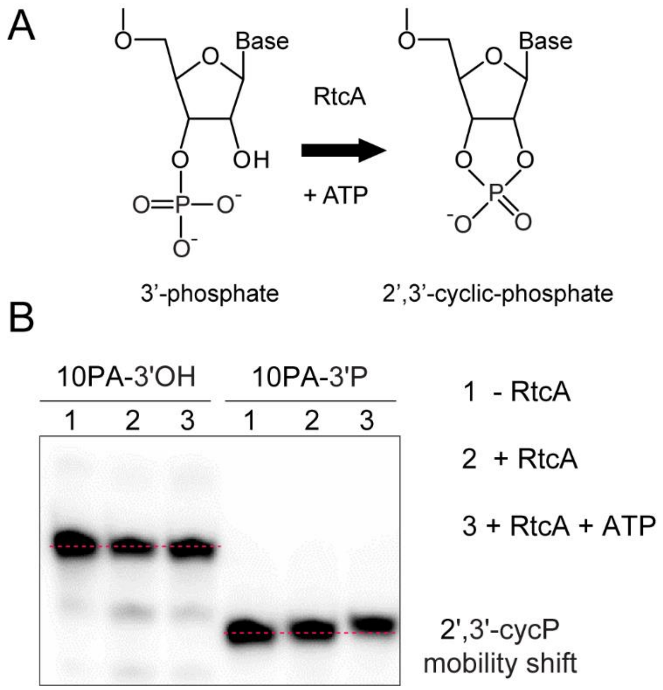


**Figure S4. RtcA catalyzes the conversion of 3’-phosphate to 2’,3’-cyclic phosphate.** (**A**) Structure of 3’-phosphate and its subsequent conversion to 2’,3’-cyclic phosphate catalyzed by RtcA. The reaction requires ATP. (**B**) Analysis of reaction products by denaturing PAGE when either a 3’-OH or 3’-phosphate terminated RNA was used as substrate with RtcA in the absence or presence of ATP. The conversion of the 3’-phosphate to 2’,3’-cyclic phosphate results in a slight mobility shift of the $^{32}$P-labeled RNA. This only occurs in the presence of ATP. The horizontal red dashed line is drawn to allow better visualization of the upward mobility shift of the 2’,3’-cyclic phosphate terminated RNA compared to the 3’-phosphate terminated RNA. No change in mobility was observed with a 3’-OH terminated RNA.

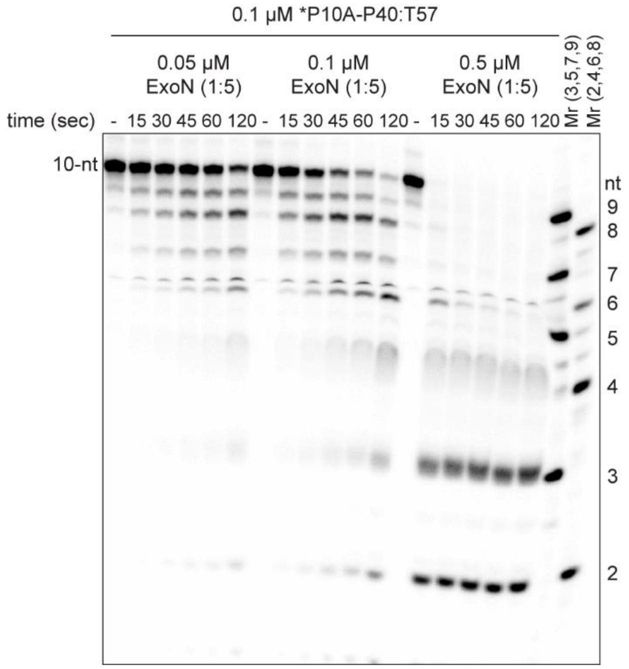


**Figure S5. ExoN appears processive under conditions of enzyme excess; however, ExoN is actually distributive.** Analysis of reaction products by denaturing PAGE from using the *P10A-P40:T57 dsRNA substrate. Reactions contained 0.1 μM RNA and either 0.05, 0.1 or 0.5 μM ExoN (1:5) and were quenched at the indicated times.

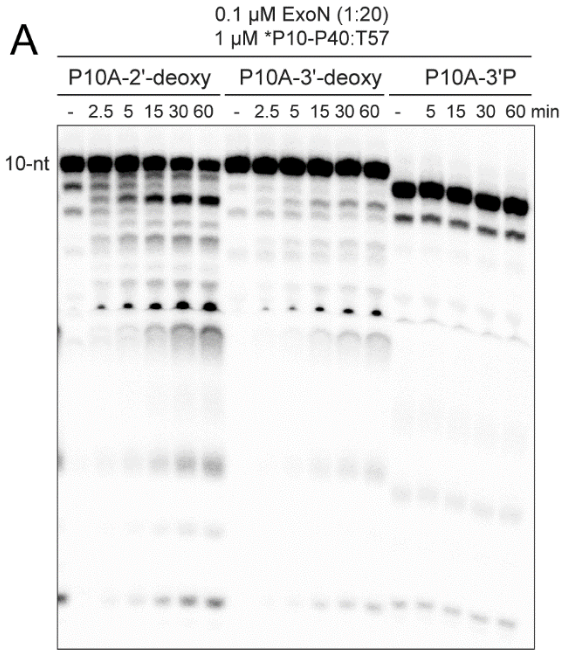


**Figure S6. Cleavage of various 3’-terminated (2’-deoxy, 3’-deoxy, 3’-phosphate) RNA substrates.** (**A**) Analysis of reaction products from reactions with using 2’-deoxy, 3’-deoxy or 3’-phosphate terminated RNA when RNA is in excess of ExoN. Reactions contained 0.1 µM ExoN (1:20) and 1 µM *10P-P40:T57 dsRNA substrate and were quenched at the indicated times. (**B**) Analysis of reaction products from reactions with using 2’-deoxy, 3-’deoxy or 3’-phosphate terminated RNA when ExoN is in excess of RNA. Reactions contained 0.1 µM ExoN (1:20) and 0.05 µM *10P-P40:T57 dsRNA substrate and were quenched at the indicated times. In each case, the primer used was *10PA-2’d, *10PA-3’d, or 10PA-3’P.

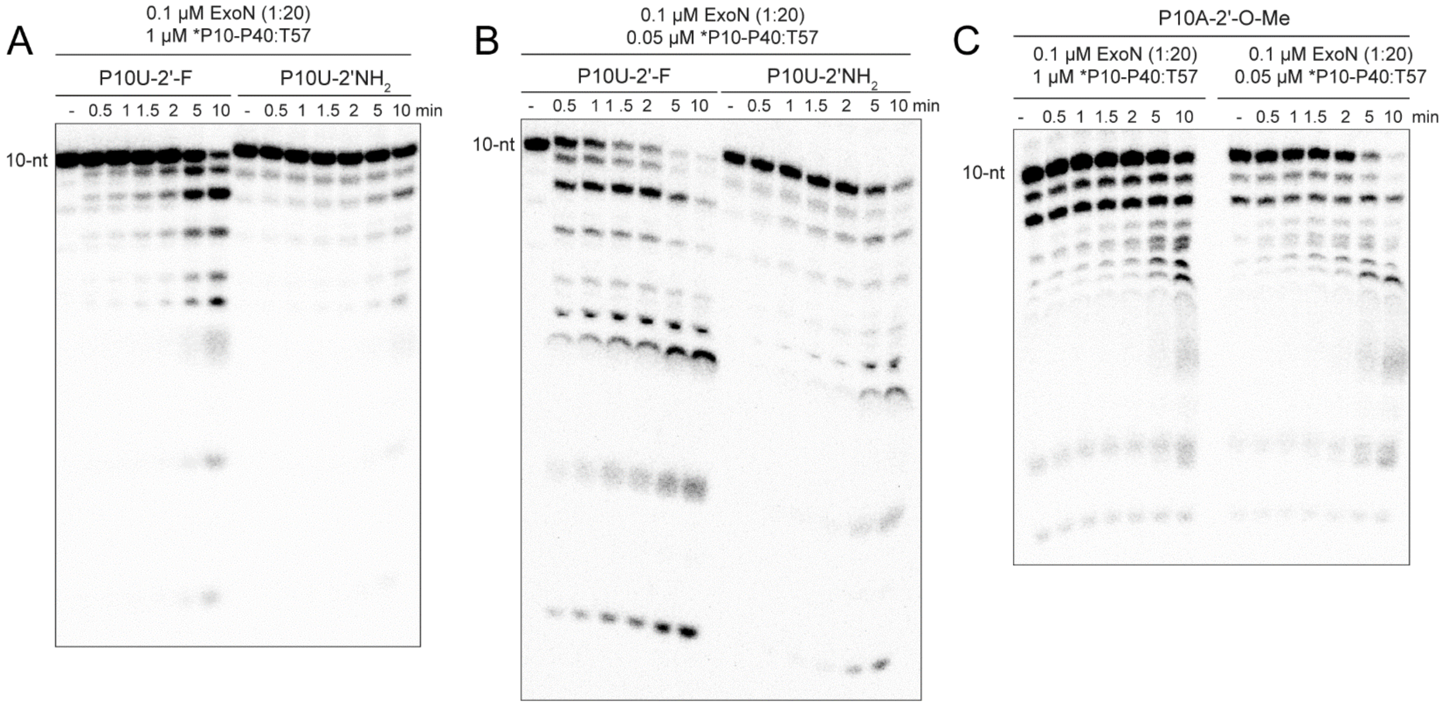


**Figure S7. Cleavage of various 3'-terminated (2'-F, 2'-NH2, 2'-OMe) RNA substrates.** (**A**) Analysis of reaction products from reactions with using 2'-fluoro, 3'-amino or 2'-O-methyl terminated RNA when RNA is in excess of ExoN. Reactions contained 0.1 µM ExoN (1:20) and 1 µM *10P-P40:T57 dsRNA substrate and were quenched at the indicated times. In each case the primer used was *10PU-2'F, *10PU-2'NH2, or 10PA-2'-O-Me. (**B**) Analysis of reaction products from reactions with using 2'deoxy, 3'deoxy or 3'phosphate terminated RNA when ExoN is in excess of RNA. Reactions contained 0.1 µM ExoN (1:20) and 0.05 µM *10P-P40:T57 dsRNA substrate and were quenched at the indicated times. In each case the primer used was *10PU-2'F, *10PU-2'NH2, or 10PA-2'-O-Me.
